# Enzyme-constrained models and omics analysis of Streptomyces coelicolor reveal metabolic changes that enhance heterologous production

**DOI:** 10.1101/796722

**Authors:** Snorre Sulheim, Tjaša Kumelj, Dino van Dissel, Ali Salehzadeh-Yazdi, Chao Du, Gilles P. van Wezel, Kay Nieselt, Eivind Almaas, Alexander Wentzel, Eduard J Kerkhoven

## Abstract

Many biosynthetic gene clusters (BGCs) require heterologous expression to realize their genetic potential, including silent and metagenomic BGCs. Although the engineered *Streptomyces coelicolor* M1152 is a widely used host for heterologous expression of BGCs, a systemic understanding of how its genetic modifications affect the metabolism is lacking and limiting further development. We performed a comparative analysis of M1152 and its ancestor M145, connecting information from proteomics, transcriptomics, and cultivation data into a comprehensive picture of the metabolic differences between these strains. Instrumental to this comparison was the application of an improved consensus genome-scale metabolic model (GEM) of *S. coelicolor*. Although many metabolic patterns are retained in M1152, we find that this strain suffers from oxidative stress, possibly caused by increased oxidative metabolism. Furthermore, precursor availability is likely not limiting polyketide production, implying that other strategies could be beneficial for further development of *S. coelicolor* for heterologous production of novel compounds.

## Introduction

The bacterium *Streptomyces coelicolor* has been the *de facto* model actinomycete for the production of antibiotics. Being known for over 100 years, the interest in this organism predates the golden age of antibiotic research. With its complex life cycle, featuring mycelial growth and differentiation, spore formation, programmed cell death and the ability to produce multiple coloured secondary metabolites it has assisted greatly in our understanding of how streptomycetes sense their surrounding (Hahn, Oh, and Roe 2002; Hutchings et al. 2004; Nothaft et al. 2010, 2; Rigali et al. 2008; Sola-Landa et al. 2005), activate their developmental cycle (Chandra and Chater 2014) and regulate the production of antibiotics (Nieselt et al. 2010; Thomas et al. 2012). Further aided by the publication of its genome sequence (Bentley et al. 2002), the antibiotic coelimycin P1 (yellow), produced from the formerly cryptic polyketide gene cluster known as *cpk*, was added to this list (Gomez-Escribano et al. 2012). Today, the widespread use of *S. coelicolor* continues as a host for heterologous production of biosynthetic gene clusters (BGCs) (Castro et al. 2015; Gomez-Escribano and Bibb 2011; 2014; Kumelj et al. 2018; Thanapipatsiri et al. 2015; Yin et al. 2015). Heterologous expression is a powerful strategy for novel compound discovery from BGCs that are either natively silent or originate from an unculturable source (Nepal and Wang 2019). These BGCs represent an untapped resource of microbial biodiversity, nowadays made evident and accessible due to recent advances within the fields of metagenomics, molecular biology and bioinformatics (Rutledge and Challis 2015).

The efficiency of *S. coelicolor* as a heterologous production host relies on a metabolism that has evolved to provide the necessary precursors to produce a broad range of complex molecules. Many of these molecules are produced when the strain is experiencing nutrient-limiting conditions that lead to growth cessation and complex re-modelling of its metabolism (Wentzel, Bruheim, et al. 2012). Metabolic switching in response to phosphate and glutamate depletion has been studied in detail at a variety of metabolic levels in *S. coelicolor* M145 (Nieselt et al. 2010; Thomas et al. 2012; Wentzel, Sletta, et al. 2012), a the most well-known wild-type strain devoid of the two plasmids SCP1 and SCP2 present in the parent strain *S. coelicolor* A3(2) (Kieser et al. 2000). This has unravelled a complex sequence of switching events that ultimately lead to the biosynthesis of calcium-dependent antibiotic (CDA), and the coloured antibiotics actinorhodin (Act, blue) and undecylprodigiosin (Red, red). The biosynthesis of coelimycin P1 occurs earlier than the three other compounds in the growth cycle and appears to be independent of the major metabolic switch (Nieselt et al. 2010).

To improve *S. coelicolor* M145 as a host for heterologous BGCs expression, strain M1146 was created by the sequential deletion of its four major BGCs (*act, red, cda* and *cpk*) (Gomez-Escribano and Bibb 2011). This should increase precursor availability for the production of a whole range of heterologous products and provides a cleaner chromatographic background to more easily identify novel compounds. *S. coelicolor* M1152 is a derivative of M1146, that besides the deletion of the four main BGCs bears the C1298T point mutation in the *rpoB* gene that encodes the beta subunit of RNA polymerase. This mutation was shown to have strong positive effects on the production of various antibiotics (Gomez-Escribano and Bibb 2011; Hu, Zhang, and Ochi 2002). Up to now, M1152 is a preferred general ‘superhost’ for heterologous BGC expression (Braesel, Tran, and Eustáquio 2019; Castro et al. 2015; Kepplinger et al. 2018; T. Li et al. 2013; Thanapipatsiri et al. 2015) and is the starting point for further strain development.

Previous research on the metabolism of *S. coelicolor* M1152 have been confined to transcriptome profiling of batch fermentations (Battke, Symons, and Nieselt 2010; Jäger, Battke, and Nieselt 2011; Liao, Smyth, and Shi 2014; Love, Huber, and Anders 2014; Mi et al. 2019), and further development of this strain as a ‘superhost’ calls for a better understanding of how the genetic modifications have affected M1152s regulatory system and metabolism. To this end we measure both protein and transcript levels of both M1152 and its parent strain, M145, at different time steps during batch fermentation where the metabolic switch is triggered by depletion of phosphate. Since enzymes are catalysing most metabolic transformations, assessing protein abundance provides information about the metabolic capacity of the organism. Furthermore, we do not only consider the protein abundances in isolation, but use these measurements to confine fluxes predicted by a genome-scale metabolic model (GEM) of *S. coelicolor* to the maximum capacity of the enzymes. By doing so we propagate differences in the abundance of individual enzymes in M145 and M1152 to metabolic rearrangements on the systems level.

The metabolic network in the cell is described in a GEM (Gu et al. 2019). GEMs are both valuable resources of strain-specific knowledge, mathematical models able to predict steady-state flux distributions, and frameworks for interpretation and integration of different ‘omics’ data, e.g. transcriptomics and proteomics (Robinson and Nielsen 2016). The increased interest in using genome-scale models of *S. coelicolor* is conspicuous. Since the first reconstruction in 2005 (Borodina, Krabben, and Nielsen 2005), five GEMs have been published (Alam et al. 2010; Amara, Takano, and Breitling 2018; M. W. Kim et al. 2014; Kumelj et al. 2018; Wang et al. 2018), including three in 2018: iKS1317 (Kumelj et al. 2018), Sco4 (Wang et al. 2018) and iAA1259 (Amara, Takano, and Breitling 2018). Additionally, as a model organism for the *Actinomycetes,* the GEMs of *S. coelicolor* are frequently used as template for model development of closely related strains (Mohite et al. 2019), such as *S. clavuligerus* (Toro et al. 2018), *Saccharopolyspora erythraea* (Licona-Cassani et al. 2012) and *S. lividans* (Valverde, Gullón, and Mellado 2018). The recent updates of the *S. coelicolor* GEM were developed in parallel by different research groups: while all groups share the common interest of utilizing a high-quality model for predictions and data analysis, the prevailing approach of independent parallel development is inefficient. Additional to duplicating a considerable amount of work, lack of common standards for documentation of progress and issues, evaluation of model performance, as well as the use of different annotations makes it cumbersome to compare and merge models.

To increase the rate and quality of model reconstruction, in this study two research groups of the *S. coelicolor* GEM community, responsible for two of the latest model updates (Kumelj et al. 2018; Wang et al. 2018), have joined forces to merge existing GEMs of *S. coelicolor* into one consensus-model that is publicly hosted on GitHub and can be continuously updated and improved by all members of the community. Hosting the model on GitHub has many advantages: (i) open access and contribution; (ii) version control; (iii) continuous development and integrated quality control with memote (Lieven et al. 2018); (iv) new improvements released instantly (no publication lag time); and (v) complete documentation of model reconstruction. Such an approach has historic precedents: model reconstruction as a community effort has been a success for the human GEM (Thiele et al. 2013), baker’s yeast (Aung, Henry, and Walker 2013; Dobson et al. 2010; Heavner et al. 2012; 2013; Herrgård et al. 2008; Lu et al. 2019) and Chinese Hamster Ovary cells (Hefzi et al. 2016). The recent developments in *S. coelicolor* model and strain improvements in different research groups prove that it is an opportune time now to join forces in the *Streptomyces* modelling efforts as well.

## Results

### Reconstruction of the consensus genome-scale model of *S. coelicolor*

We conducted a stepwise reconstruction of Sco-GEM, the consensus genome-scale metabolic model of *S. coelicolor*, while tracking development using Git for version control (**Figure 1A**; **Data Set S1, Tab 1**). Sco-GEM is the most comprehensive and highest quality GEM of this organism (**Figure 1B**), comprising 1777 genes, 2612 reactions, 2073 metabolites and a memote score of 77%, which is indicative of the overall model quality (Lieven et al. 2018). Sco-GEM features an accuracy of 96.5% and 74.5% (**Figure 1C**) in predicting correct phenotypes for growth environments and knockout mutants, respectively, yielding in total a Matthews Coefficient of Correlation of 0.53 with the test data previously described (Kumelj et al. 2018).

**Figure 1:**
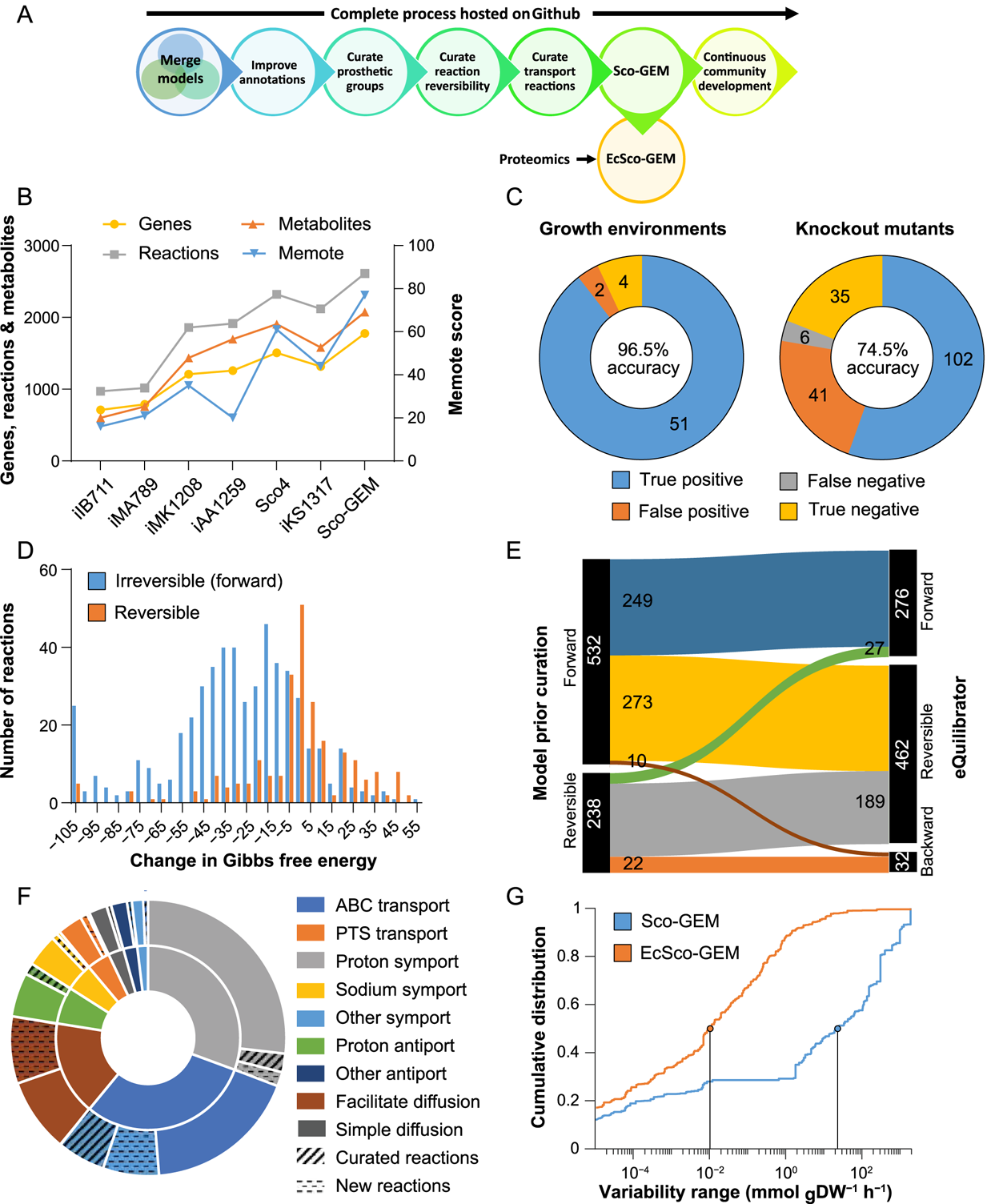
Sco-GEM development and analysis. A) Schematic overview of the various steps in the Sco-GEM reconstruction process. B) The overall memote score and number of genes, reactions and metabolites for the 7 published *S. coelicolor* GEMs. C) Assessment of the model quality by comparing *in vivo* observations with *in silico* predictions across in total 241 tests: accuracy = 0.80; sensitivity = 0.96; specificity = 0.48; Matthews Correlation Coefficient = 0.53. D) The change in Gibbs free energy for 770 reactions that were annotated as either reversible or forward (i.e. forward irreversible) in the model prior curation of reaction reversibility. The histogram is truncated at −105 kJ/mol, and more negative values are assigned to the leftmost bin. E) Analysis and comparison of the directionality and reversibility of reactions prior curation and the direction inferred from the change in Gibbs free energy as estimated by eQuilibrator. Reactions labelled “forward” or “backward” are irreversible. F) Overview of the 369 transport reactions included in Sco-GEM, whereof 42 were curated and 65 added during this work. The inner ring categorizes the reactions into 9 different subgroups, while the outer ring displays the amount of curated and added reactions within each category. In the outer ring, the sections representing curated and new reactions are hatched and dotted, respectively. G) Comparison of cumulative flux variability distributions in Sco-GEM and EcSco-GEM.

With the recently published iKS1317 model (Kumelj et al. 2018) as a starting point, Sco-GEM was first developed by including genes, reactions and metabolites from the equally recently published models iAA1259 (Amara, Takano, and Breitling 2018) and Sco4 (Wang et al. 2018). While the curations from iAA1259 were primarily related to coelimycin P1, butyrolactone, xylan and cellulose pathways, the 377 reactions added to Sco-GEM from Sco4 were scattered across a large range of different subsystems, covering both primary and secondary metabolism (**Figure S1**). Subsequent to merging the existing *S. coelicolor* GEMs, we performed a number of further curations of the model (**Figure 1A**): including improvement of annotations, both in terms of coverage and number of different databases, e.g. KEGG (M. Kanehisa 2000; Minoru Kanehisa et al. 2019), BioCyC (Karp et al. 2017), ChEBI (Hastings et al. 2016) and MetaNetX (Moretti et al. 2016). All reactions and metabolites have been given identifiers according to the BiGG namespace (King et al. 2016), and all reactions are categorized into 15 different subsystems, covering 128 different pathways.

The biomass composition was curated to reflect estimated levels of prosthetic groups that are associated to cellular proteins. Proteomics data, as discussed below, were used to estimate protein levels, while UniProt (The UniProt Consortium 2019) provided annotations of proteins with prosthetic groups, which was used to estimate overall prosthetic group levels (**Data Set S1, Tab 2**).

### Reaction reversibility updated for almost a third of queried reactions

The determination of reaction directionality and reversibility is an important step in a GEM reconstruction (Thiele and Palsson 2010). However, the thermodynamic consistency of reactions was not considered in previous *S. coelicolor* models. We calculated Gibbs free energy changes for 770 of the 2612 model reactions (**Data Set S1, Tab 3**) using eQuilibrator (Flamholz et al. 2012), and found hardly any consistency between the calculated change in Gibbs free energy and the reversibility previously assigned to the model reactions (**Figure 1D**). To address this issue we decided to reassign the reversibility of the model reactions by using a relatively lenient threshold of −30 kJ/mol to classify a reaction as irreversible (Bar-Even et al. 2012; Feist et al. 2007); with the intent not to over-constrain the model (**Figure 1E**). The proposed changes in reversibility were evaluated against growth and knockout data (Kumelj et al. 2018), discarding 61 of the 332 proposed reactions, and consequentially, the flux bounds of 271 reactions were modified (see Transparent Methods). Additionally, all ATP-driven reactions were manually curated and generally assumed irreversible unless they had an estimated positive change in Gibbs free energy or were known to be reversible. Examples of this include nucleoside diphosphate kinase (Chakrabarty 1998) and ATP synthase (Yoshida, Muneyuki, and Hisabori 2001). The manual curation of ATP-driven reactions led to a change in reversibility for 56 reactions.

### Curation of transport reactions

As transport reactions have previously not been extensively curated in *S. coelicolor* models, we performed a thorough curation of transporters by querying various databases and BLAST analysis as detailed in Materials and Methods. This culminated in adding 43 new transport reactions and updating 39 of the 262 existing reactions in Sco-GEM (**Figure 1F**; **Data Set S1, Tab 4**). The majority of the transporters comprises primary active transport proteins and secondary carriers (46%), in accordance with previous work (Getsin et al. 2013). Most primary active transporters are ATP-binding cassette (ABC) transporters (30%), while proton symports (30%) dominate the secondary carriers.

### Development of the enzyme-constrained model EcSco-GEM

To include explicit constraints regarding enzymes catalysing metabolic reactions, the GECKO formalism (Sánchez et al. 2017) was applied to consider that catalysing capacity is constrained by enzyme turnover rates (kcat) and abundances. The GECKO toolbox modifies the structure of an existing GEM to integrate turnover rates and proteome data.

Consequentially, this constrains the range of estimated fluxes to a biologically feasible range as determined by the amount and efficiency of each enzyme. Note that this approach regards the maximum catalytic activities but does not consider other kinetic parameters such as affinity constants. The overall flux variability of the resulting enzyme-constrained model (EcSco-GEM) is drastically reduced compared to the classic genome-scale model (**Figure 1G**), particularly due to the considerably reduced fraction of reactions that have very high (10^1^) flux variability. As reactions with high variability result in low certainty in the estimated fluxes, the observed reduction in flux variability is therefore a qualitative measure of the increased accuracy achieved by constraining the range of possible fluxes to those satisfying the limitation in protein allocation.

In our endeavour to describe the metabolic differences between M145 and M1152 we generated in total 17 time- and strain-specific enzyme-constrained models by combining EcSco-GEM with estimated growth-, secretion- and uptake rates, as well as proteome data from cultivations that are detailed and analysed below.

### Framework for further development of Sco-GEM by the community

The Sco-GEM model is hosted as an open repository as suggested by memote, a recently developed tool for transparent and collaborative model development (Lieven et al. 2018). The memote tool is incorporated in the repository through Travis CI and tracks the model development on every change of the model. Sco-GEM v1.2.0 achieved a memote-score of 77%, which is superior to any previous model of *S. coelicolor* (**Figure 1B**; **Supplemental Information**).

Hosting Sco-GEM on GitHub with memote integration ensures continuous quality control and enables public insight into all aspects of model reconstruction and curation: any user can report errors or suggest changes through issues and pull requests. As contributions to the model development are fully trackable and can therefore be credited fairly, Sco-GEM is positioned as a community model that we envision to be continuously updated and widely used by the *S. coelicolor* research community. While the major steps of model reconstruction have been detailed in the preceding sections, every detail of the process and every iteration of the model is accessible on the public model repository at https://github.com/SysBioChalmers/Sco-GEM.

In the remaining parts of the Results section, we have applied Sco-GEM along with transcriptome and proteome data, to study and compare the responses of *S. coelicolor* M145 and M1152 to phosphate depletion on a systems level and for the first time provide detailed insight into the distinct physiological features of engineered ‘superhost’ strain M1152, which will be of value for its further development.

### Random sampling of enzyme-constrained GEMs capture metabolic rearrangements in response to phosphate depletion in M145

To evaluate whether the (Ec)Sco-GEM models can simulate behaviours of *S. coelicolor* metabolism, we analysed time-course sampled cultivations of secondary metabolite producing strain M145 using the generated models. For that purpose, *S. coelicolor* M145 was cultivated in batch fermentations using standardized protocols reported earlier (Wentzel, Bruheim, et al. 2012). Cultures were sampled for ‘omics data, as well as substrate utilization and secondary metabolite measurements to identify regulatory, proteomic and metabolic changes during the metabolic switch. The online and offline measurements showed that phosphate depletion in the cultivation medium was reached approximately 35 hours after inoculation. Shortly after, the culture growth ceased, and first Red and subsequently Act were detected in the culture medium (**Figure 2A and 2B**). Act levels were determined by measuring the amount of total blue pigments (TBP) because this covers both the intracellular and secreted variants of actinorhodin, and is considered to be the preferred method (Wentzel, Bruheim, et al. 2012; Bystrykh et al. 1996). Both D-glucose and L-glutamate were consumed concomitantly, and their consumption continued after phosphate depletion, while both remained in excess until the end of cultivation. Note that *Streptomyces* can utilize intracellular phosphate storages after the medium is phosphate depleted (Smirnov et al. 2015). The RNA-seq and untargeted proteomic data were analysed in the light of previous studies (Nieselt et al. 2010; Thomas et al. 2012) and were in good agreement with data previously obtained from microarrays or targeted proteomics (Alam et al. 2010; Nieselt et al. 2010) (**Figure 2C** and **S2**). This confirmed the high reproducibility of the experiments across independent cultivations and high reliability of the chosen cultivation and analytic procedures (**Figure 2**).

**Figure 2:**
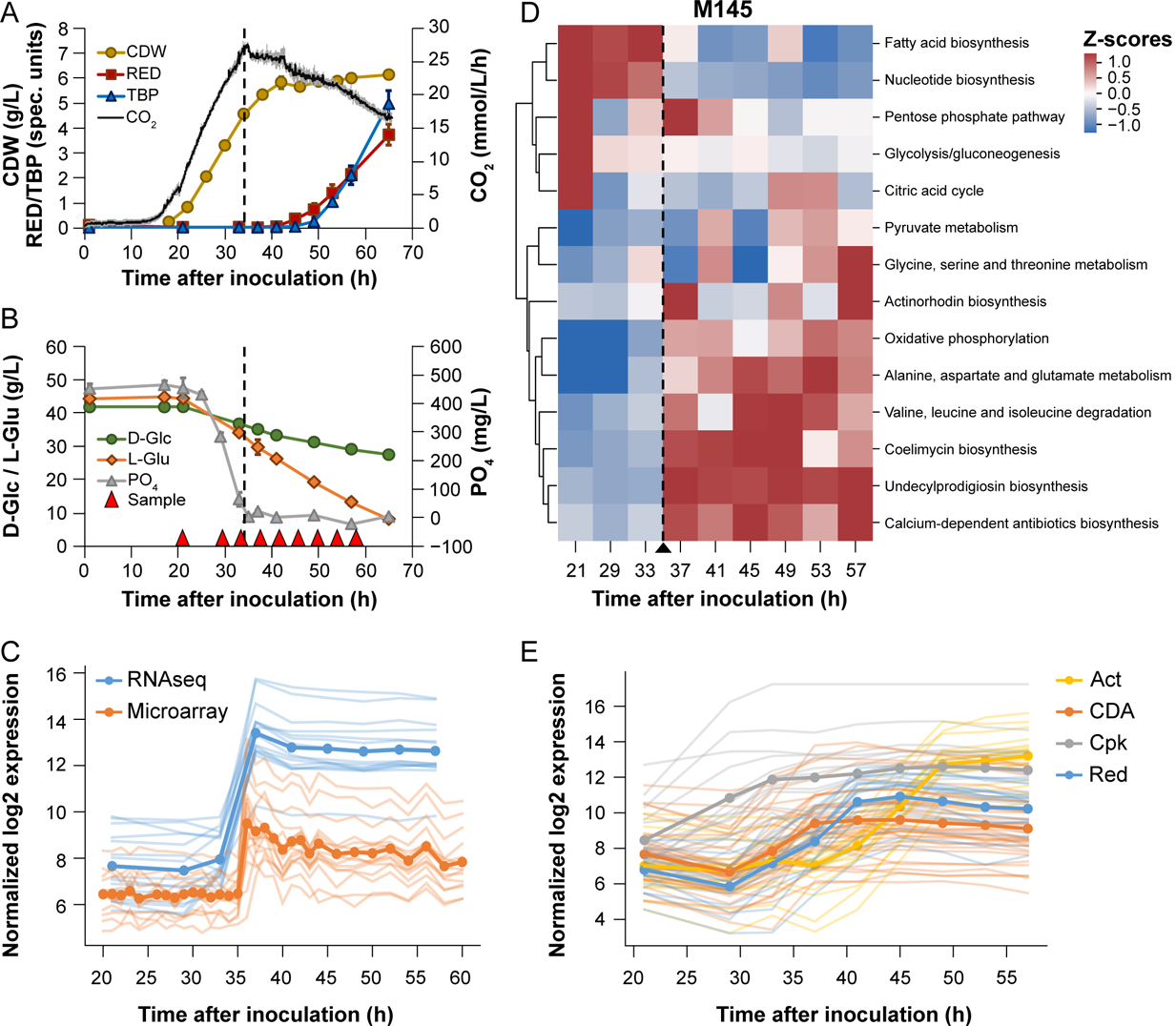
Batch cultivation of *S. coelicolor* M145 and the effect of phosphate depletion. Compounds produced (A) and consumed (B) during batch fermentation of *S. coelicolor* M145. Time points for sampling for transcriptome and proteome analysis are indicated with red triangles. The dashed vertical line indicates when phosphate in the medium has been depleted. Error bars are standard deviations of three biological replicates. CDW, Cell Dry Weight; Red, undecylprodigiosin; TBP, Total Blue Pigments/actinorhodins; CO2 volume corrected respiration; D-Glc, D-glucose; L-Glu, L-glutamate; PO4, phosphate. C) Comparison of previously published microarray data (Nieselt et al. 2010) and RNA-seq data (this study) for genes previously found to respond to phosphate depletion (Nieselt et al. 2010). The transparent lines correspond to individual genes, while the bold lines represent the average expression level for each data set. D) Clustered heatmap of CO2-normalized Z-scores for each of the top 10 varying pathways plus the pathways for the 4 major BGCs in M145, as revealed by simulations with the proteomics-integrated EcSco-GEM model. The pathways are sorted based on hierarchical clustering to facilitate visual interpretation of similarity between pathways. The dashed vertical line indicates the time point of the metabolic switch. E) RNA-seq data of the 4 major BGCs show the onset of biosynthesis of actinorhodin (Act), calcium-dependent antibiotic (CDA), coelimycin P1 (Cpk) and undecylprodigiosin (Red) at different time points during the batch fermentations of M145.

The proteome data were incorporated into EcSco-GEM to yield time-specific metabolic models of M145, giving insight on the changes occurring in the metabolic activity of different pathways during batch cultivation. Metabolic fluxes were estimated using an unbiased approach of random sampling, as alternative to optimization of a well-defined cellular objective used in flux balance analysis (Orth, Thiele, and Palsson 2010). It is possible that *S. coelicolor* is wired to maximize its growth rate prior to phosphate depletion, but after the metabolic switch, it is difficult to define a clear cellular objective. We applied an approach that samples the vertices of the solution space (Bordel, Agren, and Nielsen 2010), and used their mean values to compare the metabolic fluxes between the two strains and between different time points. The variation in predicted fluxes through different pathways in M145 is an initial validation of the approach (**Figure 2D**): the most drastic change in fluxes occur in response to phosphate depletion, in agreement with observations in the transcriptome, metabolome and proteome (Nieselt et al. 2010; Thomas et al. 2012; Wentzel, Sletta, et al. 2012).

The response to phosphate depletion from the medium is achieved by a set of genes, positively regulated by PhoP, that are involved in phosphate scavenging, uptake and saving (Martín et al. 2012; Martín-Martín et al. 2018; Sola-Landa, Moura, and Martín 2003). In our cultivations the metabolic switch can be readily identified from the RNA-seq data by the rapid upregulation of this regulon after 35 hours of cultivation in M145 (**Figure 2C**), thereby corroborating the model simulations (**Figure 2D**) and providing a more detailed picture of the underlying regulation. PhoP also represses nitrogen assimilation (Martín, Rodríguez-García, and Liras 2017), which can partly explain the change in amino acids metabolism after phosphate depletion (**Figure 2D**). Indeed, from the RNA-seq data we find that glutamate import, the glutamate sensing system *gluR-gluK* (L. Li, Jiang, and Lu 2017), *glnR* (Fink et al. 2002) and *glnA* are downregulated immediately subsequent to phosphate depletion (**Figure S3**). Since PhoP is also known to regulate negatively the biosynthesis of secondary metabolites, the switching of its expression likely delays these pathways (Martín 2004; Martín, Rodríguez-García, and Liras 2017). However, after 37 hours of cultivation the upregulation of the *cda* and *red* genes was observed, whereas that of the *act* genes was initiated at 41 hours (**Figure 2E**). Production of Red and Act was measurable in the culture medium after 41 and 49 hours of cultivation, respectively (**Figure 2A**). The enzyme-constrained models predict an immediate increase in fluxes through the biosynthetic pathways for the four main compounds Act, Red, CDA and coelimycin P1 after the metabolic switch (**Figure 2D**). The onset of secondary metabolism is strongly correlated with an increase in oxidative phosphorylation and a decrease in fatty acid biosynthesis in M145.

The metabolic switch was shown to be correlated with an enhanced degradation of branched-chain amino acids (valine, leucine and isoleucine), an increase in oxidative phosphorylation and a decrease in fatty acid biosynthesis (**Figure 2D** and **S4**). An active oxidative phosphorylation relies on an active TCA cycle that generates reduced co-factors whose re-oxidation by the respiratory chain generates a proton gradient that drives ATP synthesis by the ATP synthase. The feeding of the TCA cycle requires acetyl-CoA, as well as nitrogen. Nitrogen likely originates from degradation of glutamate and branched-chain amino acids, whereas acetyl-CoA likely originates from glycolysis, as well as from the degradation of these amino acids as previously demonstrated (Stirrett, Denoya, and Westpheling 2009). Indeed, the model predicts an increased flux through citrate synthase feeding acetyl-CoA into the TCA cycle (**Figure S5A).** The predicted increase in oxidative phosphorylation is supported by the RNA-seq data showing upregulation of enzymes belonging to the respiratory chain (**Figure S5B**). This is consistent with the clear correlation previously reported between high ATP/ADP ratio, resulting from an active oxidative phosphorylation, and actinorhodin production (Esnault et al. 2017). Furthermore, the consumption of acetyl-CoA by the TCA cycle to support the oxidative metabolism logically impairs fatty acids biosynthesis (Esnault et al. 2017).

The pentose phosphate pathway provides the main redox cofactor NADPH for polyketide biosynthesis, as well as to combat oxidative stress, and its model-predicted flux increase upon initiation of polyketide synthesis (**Figure 2D**) is in agreement with previous studies (Borodina et al. 2008; Jonsbu, Christensen, and Nielsen 2001). A clear positive correlation was also noticed between the biosynthesis of alanine, aspartate and glutamate, which are precursors for CDA and/or coelimycin P1 (**Figure 2D**) and the biosynthesis of these antibiotics. Similar observations were made in the antibiotic-producing *Amycolatopsis sp.* (Gallo et al. 2010). Our EcSco-GEM model proved to be in good agreement with previously reported findings, indicating that it is able to capture *S. coelicolor* metabolic behaviour.

### Model-assisted characterization of engineered *S. coelicolor* M1152 and its responses to phosphate depletion

As detailed above, EcSco-GEM shed a new light on the metabolic switch in secondary metabolite producing strain M145. *S. coelicolor* M1152 (Gomez-Escribano and Bibb 2011) is a M145 derivative devoid of the four major BGCs and bearing a point mutation in the *rpoB* gene. A better systemic understanding of M1152 metabolism would benefit to its further development as a performing host. To do so, a comparative analysis of gene expression levels and metabolic fluxes was carried out in the strains M145 and M1152.

Batch cultivations of M1152 were performed using identical conditions and comparable sampling regimes as for M145 reported above. This enabled a direct comparison of the two strains at a systems level, revealing both expected and unexpected effects of the strains’ genetic differences (**Figure 3**). As anticipated, the products of the Cpk, CDA, Red, and Act biosynthetic pathways were undetectable in M1152 (**Figure 3A**). As previously observed (Gomez-Escribano and Bibb 2011), the growth rate of M1152 is reduced compared to M145 (0.15 h^−1^ vs 0.21 h^−1^ in the initial exponential growth phase), delaying phosphate depletion by M1152 to 47 hours after inoculation (**Figure 3B**), 12 hours after M145 (**Figure 2B**).

**Figure 3:**
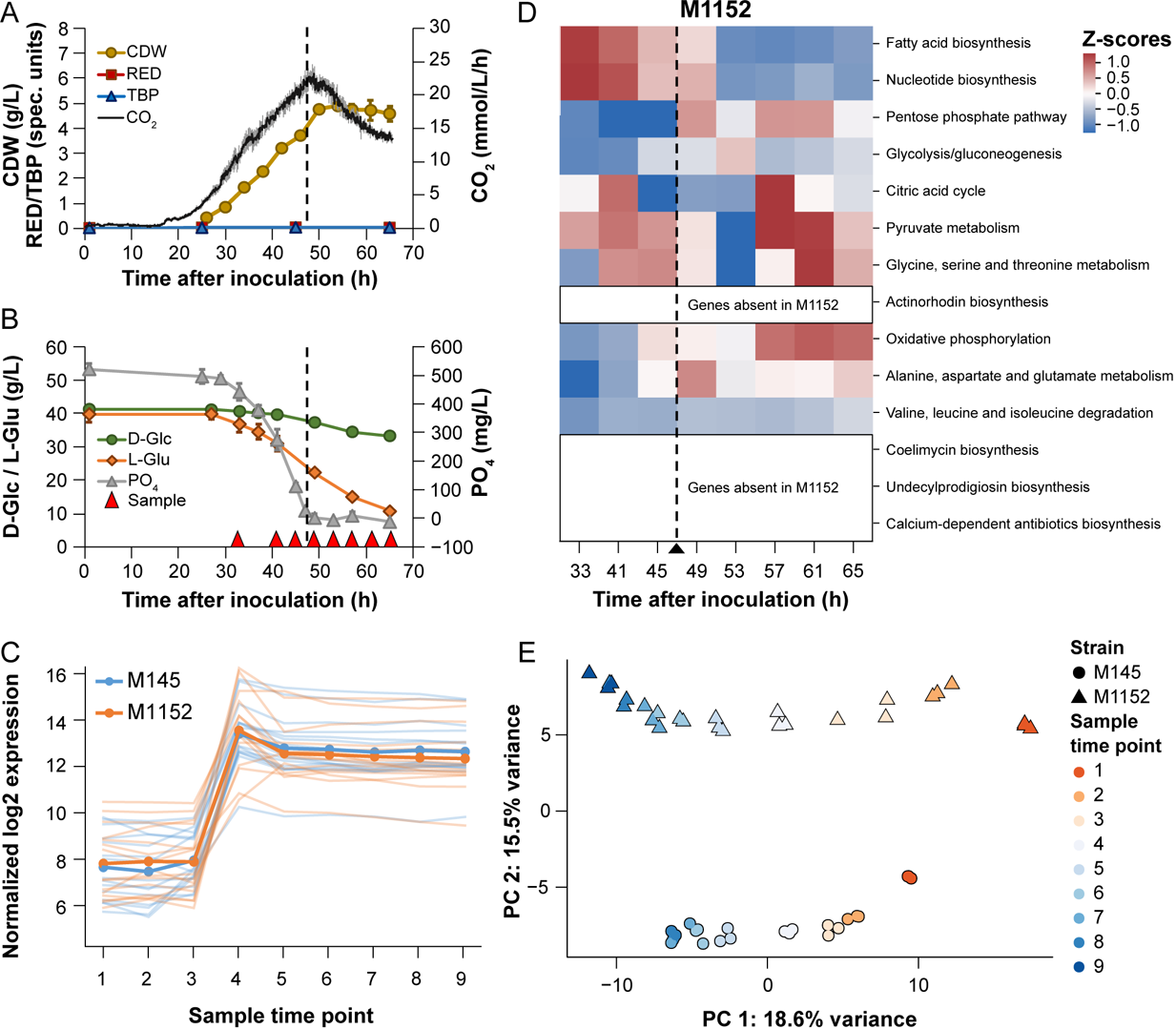
Batch cultivation of *S. coelicolor* M1152. Compounds produced (A) and consumed (B) during batch fermentation of *S. coelicolor* M1152. Time points for sampling for transcriptome and proteome analysis are indicated with red triangles. The dashed vertical line indicates when phosphate in the medium has been depleted. Error bars are standard deviations of three biological replicates. CDW, Cell Dry Weight; Red, undecylprodigiosin; TBP, Total Blue Pigments/actinorhodins; CO2, volume corrected respiration; D-Glc, D-glucose; L-Glu, L-glutamate; PO4, phosphate. C) Alignment of sample time points of M145 and M1152 cultivations based on the expression profiles of genes that were earlier found to respond to phosphate depletion in respect to the metabolic switch (Nieselt et al. 2010). D) Principle component analysis of the proteomics data for M145 (triangles) and M1152 (circles), for each time-point and culture. The first principal component separates the time points while the second principal component separates the two strains. E) CO2-normalized Z-scores of pathway fluxes predicted by EcSco-GEM for 10 of the most varying pathways in M145 and M1152. To make this heatmap comparable to the results for M145 (Figure 2D), the data is standardized for both strains simultaneously and the row order is identical.

The sampling time points for proteome and transcriptome were adjusted accordingly (**Figure 3B**), enabling pairwise comparison of measurements between the two strains. Genes responsive to phosphate depletion, members of the PhoP regulon (Nieselt et al. 2010), were used to align the different sample datasets for M145 or M1152 (**Figure 3C**). Principle component analysis of the proteome data confirms high consistency between corresponding biological replicates and incremental changes between sample points for both M145 and M1152 (mainly explained by PC1: 18.6% variance, **Figure 3D**). A clear strain dependent clustering of the data (PC2: 15.5% variance) indicates global significant differences at the protein level. EcSco-GEM was subsequently used to predict metabolic changes in M1152, and interestingly we find that most patterns in M145 are retained in M1152 (**Figure 3E**): fatty acid and nucleotide biosynthesis are still downregulated after phosphate depletion, and similar trends of upregulation at later time points are observed for oxidative phosphorylation, glycine, serine and threonine, and pyruvate metabolism. It is striking that the upregulation of the branched-chain amino acid degradation and the alanine, aspartate, and glutamate metabolism seen as a response to phosphate depletion in M145 is absent in M1152.

The different glutamate and glucose consumption rates of M145 and M1152 (**Figure 4A and 4B**) resulted in substantial metabolic differences between the two strains prior to phosphate depletion. During cultivation on SSBM-P medium, where glutamate is the sole nitrogen source, glucose and glutamate are co-consumed. M1152, as M1146 (Esnault et al. 2017), has an increased growth yield on glucose compared to M145 (**Figure S6**). It thus obtains a larger share of its carbon from glutamate (**Figure 4A and 4B**) and has consequently also a higher nitrogen availability than M145. The increased nitrogen availability does however not increase the secretion of ammonium, indicating that the consumed nitrogen is directed towards growth or production of secondary metabolites. A reduced flux through glycolysis has also been reported previously for strain M1146 (Coze et al. 2013). This might be an effect of the predicted increased concentration of ATP in M1146 compared to M145, which inhibits glucose uptake and phosphofructokinase (Coze et al. 2013; Esnault et al. 2017). Since Act was proposed to act as an electron acceptor reducing the efficiency of the oxidative phosphorylation, it is suggested that the lack of Act in M1146 causes the elevated ATP levels (Esnault et al. 2017). However, we find the largest difference in glycolytic flux at early time points, prior to phosphate depletion and Act production in M145, proving that Act itself cannot explain this observation.

**Figure 4:**
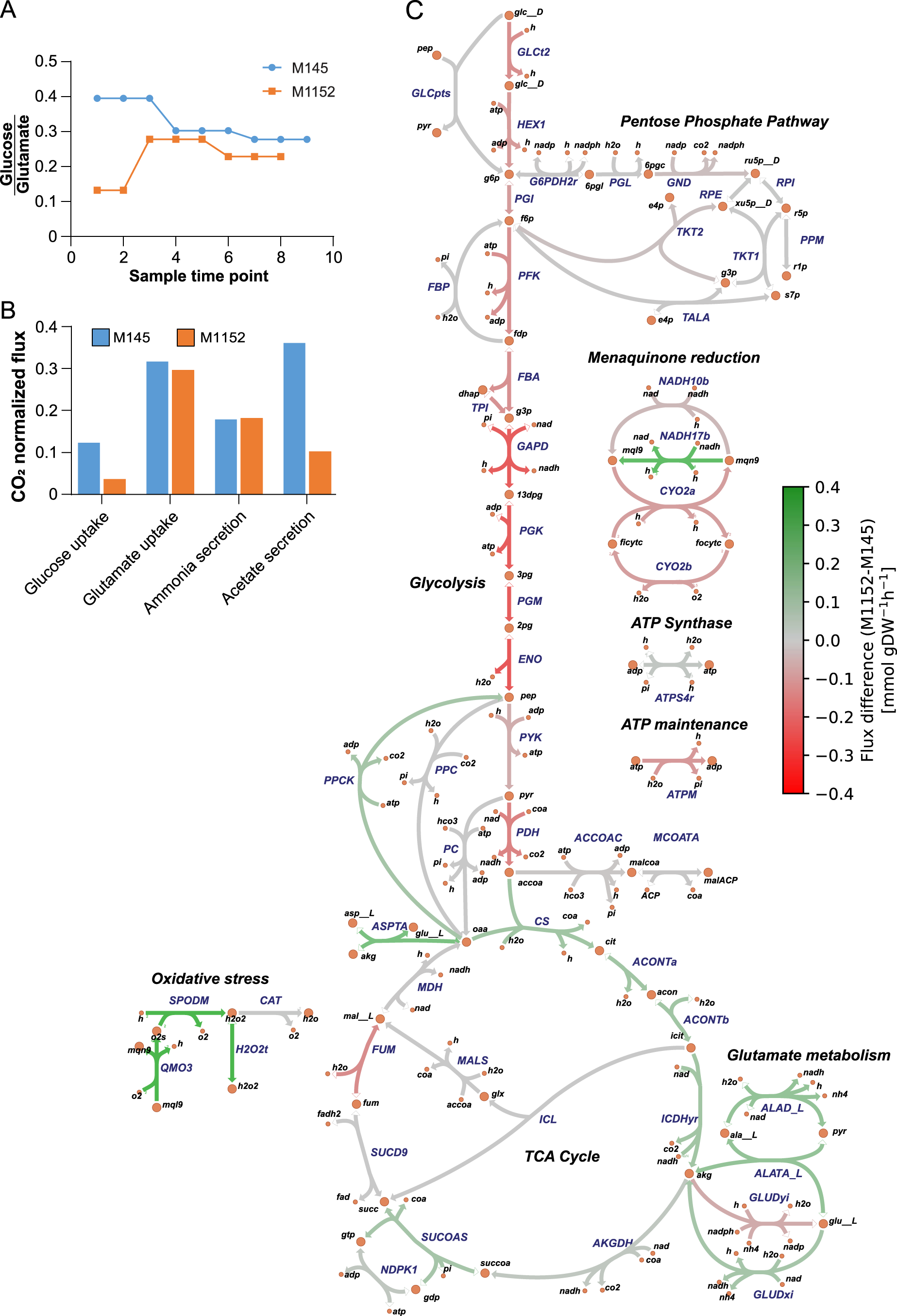
Predicted carbon fluxes in M145 and M1152. A) The ratio between estimated uptake rates of glucose and glutamate for each sample time point for M145 and M145 shows that M1152 acquires a smaller part of its carbon from glucose compared to M145. B) Bar chart showing CO2-normalized fluxes for the second sampling time point for M145 and M1152, i.e. after 29 and 41 hours, respectively. There is a clear difference in the uptake of glucose and production of acetate, while the rates are comparable for the consumption of glutamate and secretion of Ammonium. C) Comparison of predicted fluxes for the second sampling time points shows clear differences between the two strains in their relative utilization of the glycolysis and TCA cycle. The strength of the colour of the lines correspond to the flux difference between the strains; green reactions have higher flux in M1152, and red reactions have higher flux in M145.

The EcSco-GEM predicts the consequences of the reduced glucose uptake of M1152 on its central carbon metabolism, as displayed by mapping relative reaction fluxes from the second sampling time point onto a map of the central carbon metabolism in *Streptomyces* (**Figure 4C**). The map is based on the reaction network in Sco-GEM and created using Escher (King et al. 2015). A less active glycolysis in M1152 than in M145 leads to a lower carbon flow towards acetyl-CoA and thus lower excretion of acetate compared to M145 (**Figure 4B**).

Furthermore, EcSco-GEM reveals an increased flux from glutamate to alpha-ketoglutarate. Indeed, a fraction of the pool of oxaloacetate might be converted into alpha-ketoglutarate by aspartate transaminase to feed the TCA cycle. The rest might be converted into phosphoenolpyruvate (PEP) by PEP carboxykinase for gluconeogenesis because PEP carboxykinase was shown to carry higher fluxes in M1152 than in M145 (**Figure 4C**).

Since recent studies have demonstrated a negative correlation and a competition for common precursors between secondary metabolite and triacylglycerol (TAG) biosynthesis in *S. lividans* and *S. coelicolor* (Esnault et al. 2017; Millan-Oropeza et al. 2017; Craney et al. 2012), one can speculate that the acetyl-CoA/malonyl-CoA units yielded by glycolysis for the biosynthesis of antibiotics in M145 are being used for enhanced growth and/or fatty acids and TAG biosynthesis in M1152. However, this is likely not the case, as M1152 has rather a reduced growth rate compared to M145, and fatty acid biosynthesis remains downregulated after the switch (**Figure 5**). Malonyl-CoA is predominantly shuttled towards fatty acid biosynthesis through malonyl-CoA-ACP transacylase, and this consumption seems to be well-balanced by the amount of malonyl-CoA produced by acetyl-CoA carboxylase. It is noteworthy that the flux toward this acetyl-CoA/malonyl-CoA drain is 3- to 6-fold larger than the total flux going into secondary metabolite biosynthesis, even after the metabolic switch. We thus propose that together with enhanced nitrogen availability, acetyl-CoA made available from the deletion of these BGCs is used to feed the TCA cycle to support the oxidative metabolism in M1152. This would generate oxidative stress whose toxic effects might be responsible for the growth delay of this strain.

**Figure 5:**
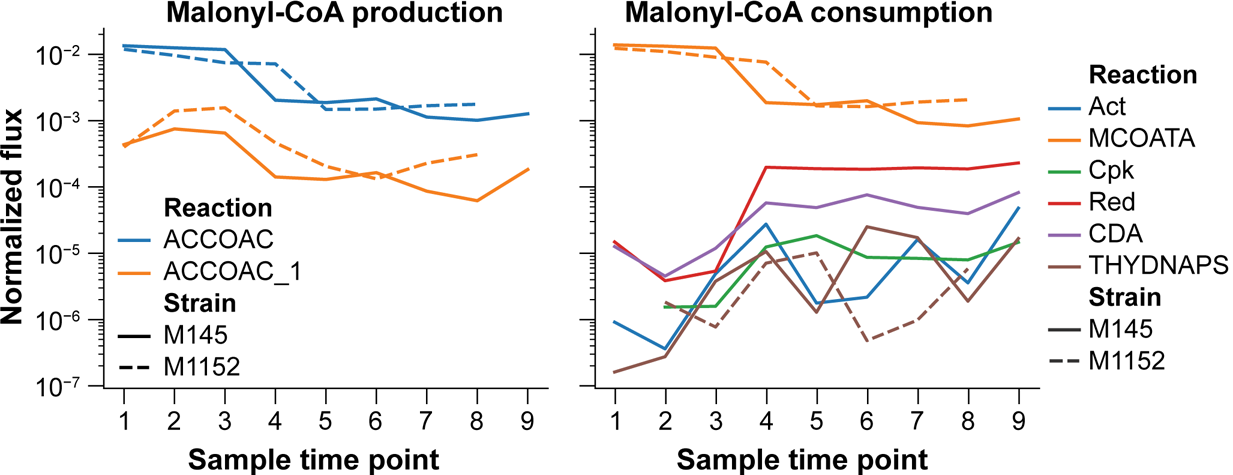
**Production and consumption of malonyl-CoA as the branching point between fatty acid biosynthesis and production of polyketides**. Both panels display CO2-normalized fluxes for both M145 and M1152 for all sampling time points as predicted by EcSco-GEM. The left panel shows the sources of malonyl-CoA, namely acetyl-CoA carboxylase (ACCOAT; blue) and acetyl-CoA carboxytransferase (ACCOAT_1; orange). We observe a downregulation of the malonyl-CoA production after the metabolic switch (between time point 3 and 4) in both strains. The right panel presents reactions consuming malonyl-CoA. The consumption is dominated by malonyl-CoA-ACP transacylase (MCOATA) leading to biosynthesis of fatty acids. The other drains for malonyl-CoA are the pathways encoded by the 4 major BGCs (Act, Cpk, Red and CDA) in addition to biflaviolin synthase (THYDNAPS).

### Transcriptome analysis reveal differential expression of global regulators

While the proteome data are an integral part of the EcSco-GEM models, RNA-seq data were used to both verify the trends and to gain further insights in the regulatory changes that are not captured by the metabolic models. As the proteomic data, the RNA-seq data showed large global differences between M1152 and M145, revealing 499 differentially expressed genes with a significance threshold of p<0.01.

Unsupervised clustering of the significantly changed genes reveal differences in regulatory systems related to redox regulation, signalling and secondary metabolism. The significantly changed genes were clustered into 7 groups with K-means clustering, with clusters 1-3 containing genes that are upregulated in M1152 compared to M145 and clusters 4-7 vice versa (**Figure S7A and Data Set S1, Tab 5**). A Gene Ontology (Ashburner et al. 2000; The Gene Ontology Consortium 2019) enrichment analysis of the seven clusters was conducted to identify upregulated processes in each of the two strains (**Figure S8**, **cf. Figure S7A**).

The enriched processes upregulated in M1152 point to increased oxidative stress (**Figure S8**): antioxidant and peroxidase activity (SCO2633 [sodF]; SCO4834-35) in addition to biosynthesis of carotenoid (SCO0185–SCO0188), a known antioxidant (Stahl and Sies 2003; Latifi, Ruiz, and Zhang 2009). The putative proteins within the cytochrome-P450 family (SCO7416–SCO7422) found in cluster 1 might also be linked to increased oxidative stress (Zangar, Davydov, and Verma 2004), but also to oxidation of precursors used for the synthesis of macrolides (Lamb et al. 2003). Indeed, by comparing the time series expression levels for genes related to oxidative stress we observe that the majority of genes related to oxidative stress are upregulated in M1152 (**Figure 6**). These changes correlate to a more active oxidative metabolism, TCA cycle and oxidative stress as predicted by Ec-ScoGEM (**Figure 4**).

**Figure 6:**
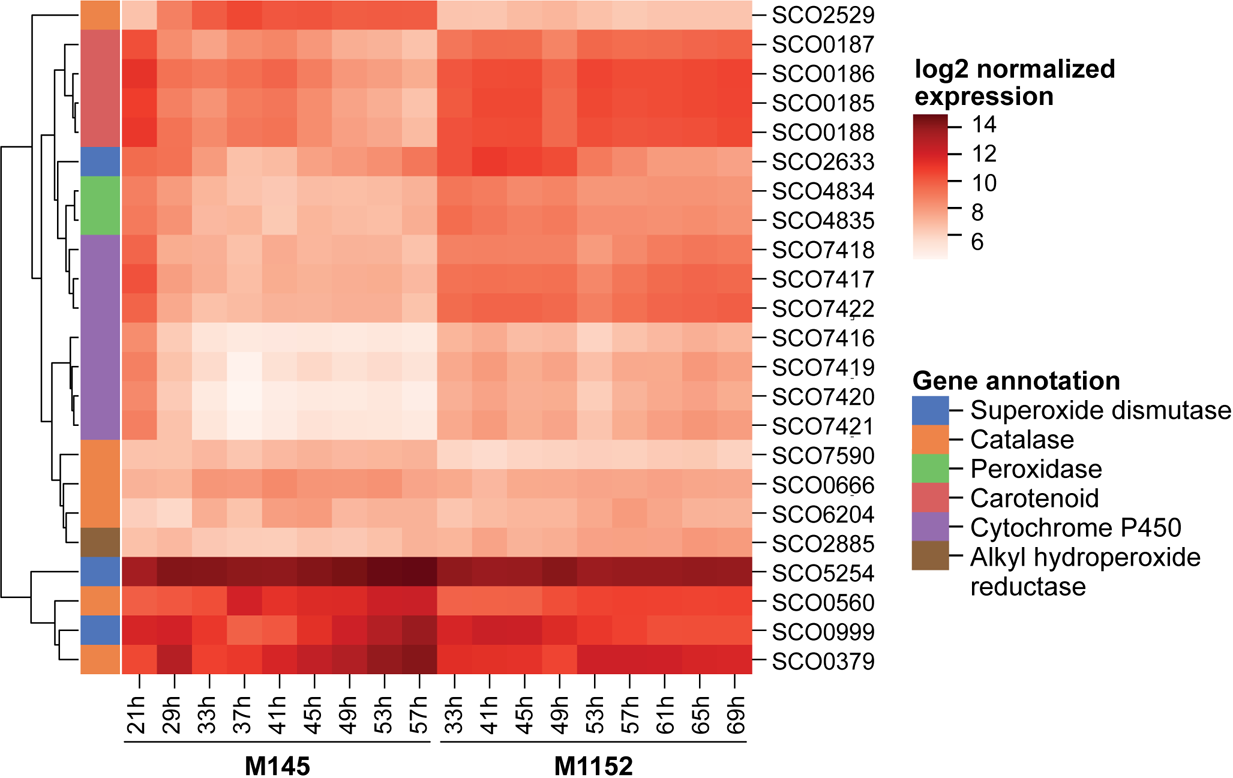
Heatmap displaying log-transformed RNA-seq data of genes associated with oxidative stress. The genes included are related to oxidative stress and either present in Sco-GEM or within the 499 differentially expressed genes. These genes are categorized based on their functional annotation to distinguish differences and similarities between these functional groups. To further enhance visual interpretatioxn the genes are ordered based on hierarchical clustering to align genes with similar expression profiles across M145 and M1152 next to each other.

In cluster 2 we find *scbA* (SCO6266) and its downstream gene *scbC* (SCO6267), which stands out by being almost 6-fold upregulated in M1152. This high expression level is likely due to the deletion of *scbR2* (SCO6286), the last gene selected to be part of the *cpk* BGC (Bednarz, Kotowska, and Pawlik 2019). Besides regulation of the *cpk* cluster, ScbR2 binds upstream of several global regulators of development and secondary metabolism, including AfsK, SigR, NagE2, AtrA, AdpA and ArgR (X. Li et al. 2015). It also acts together with ScbR to regulate ScbA, which produces the y-butyrolactone SCB1. However, when looking at the genes regulated by ScbR (X. Li et al. 2015), we only observe a clear difference in expression for genes regulated by AfsR (phosphorylated by AfsK) (Lee, Umeyama, and Horinouchi 2002; Horinouchi 2003), while this is not the case for genes regulated by ArgR, AdpA or ScbR itself (**Figure S5C-F**).

Amongst the genes upregulated in M145, in cluster 4 we find genes related to the redox regulated transcription factor SoxR (Naseer, Shapiro, and Chander 2014), and a similar pattern is observed for the entire SoxR regulon (**Figure S7B**). SoxR is known to react directly to the presence of actinorhodin (Dela Cruz et al. 2010; Shin et al. 2011), and indeed, in M145 this group of genes follows the production profile of actinorhodin, while their expression remains low in M1152 since Act is not produced. The benzoquinone Act, as electron acceptor, is thought to reduce respiration efficiency and thus energy charge as well as to combat oxidative stress (Esnault et al. 2017). Consistently, the RNA-seq data revealed that the ATP-synthase gene cluster (SCO5366–SCO5374) was upregulated almost 2-fold in M1152 compared to M145, most prominently in the stationary phase during Act production (**Figure S7C**). This agrees with observations in the M1146 strain (Coze et al. 2013). Cluster 4 also contains the genes directly up- and downstream of the deleted actinorhodin BGC in M1152 (SCO5071–SCO5072, encoding 3-hydroxyacyl-CoA dehydrogenase, and SCO5091–SCO5092, encoding a two-component flavin-dependent monooxygenase system) (Valton et al. 2008). In clusters 5, 6 and 7 we find genes with reduced expression in M1152, and the enriched processes are related to cellular and iron ion homeostasis, development, signalling and morphology. This corresponds to the delayed sporulation observed for M1152 (Gomez-Escribano and Bibb 2011).

### Elevated expression of ribosomal proteins in M1152 after phosphate depletion

An increased transcription of genes encoding ribosomal proteins could be observed in M1152 after phosphate depletion (**Figure S7D**). The *rpoB* mutation of the RNA polymerase present in M1152 is thought to induce a conformational change mimicking the binding of guanosine tetraphosphate (ppGpp) to this enzyme (Hu, Zhang, and Ochi 2002). ppGpp is synthesized in response to nutritional stress and reduces the transcription of genes related to active growth, such as genes encoding ribosomal RNAs and ribosomal proteins (Burgos et al. 2017), whereas it upregulates those involved in development/differentiation and antibiotic production (Hesketh et al. 2007; Srivatsan and Wang 2008). In consequence the upregulation of ribosomal proteins was unexpected in M1152, especially since the expression of the ppGpp regulon was not found to be significantly changed in M1152 (**Figure S5G and S5H**). We hypothesize that the ribosomal upregulation originates from the higher ATP content of M1152 compared to M145 post phosphate depletion, as high nucleoside triphosphate levels are known to have a positive impact on ribosome synthesis (Gaal et al. 1997). Such difference in ribosomal protein expression is mainly seen in the antibiotic production phase and correlated with production of Act in M145, which has a negative impact on the energetic state of the cell (Esnault et al. 2017).

### Reduced production of the polyketide germicidin in M1152

One could reasonably anticipate that the production of a secondary metabolite would increase if other drains competing for same precursor compounds were removed from the organism by gene deletion. However, the production rate of the polyketides germicidin A and B (Chemler et al. 2012), autologous to both M145 and M1152, were reduced in M1152 by 92% and 82% for germicidin A and B, respectively (**Figure 7**). This could be explained by the more active oxidative metabolism of M1152 compared to M145, as suggested by the enzyme-constrained model (**Figure 4**) and supported by the upregulation of genes associated with oxidative stress (**Figure 6**). In M1152 the pool of acetyl-CoA rather feeds the TCA cycle instead of being directed towards germicidin biosynthesis.

**Figure 7:**
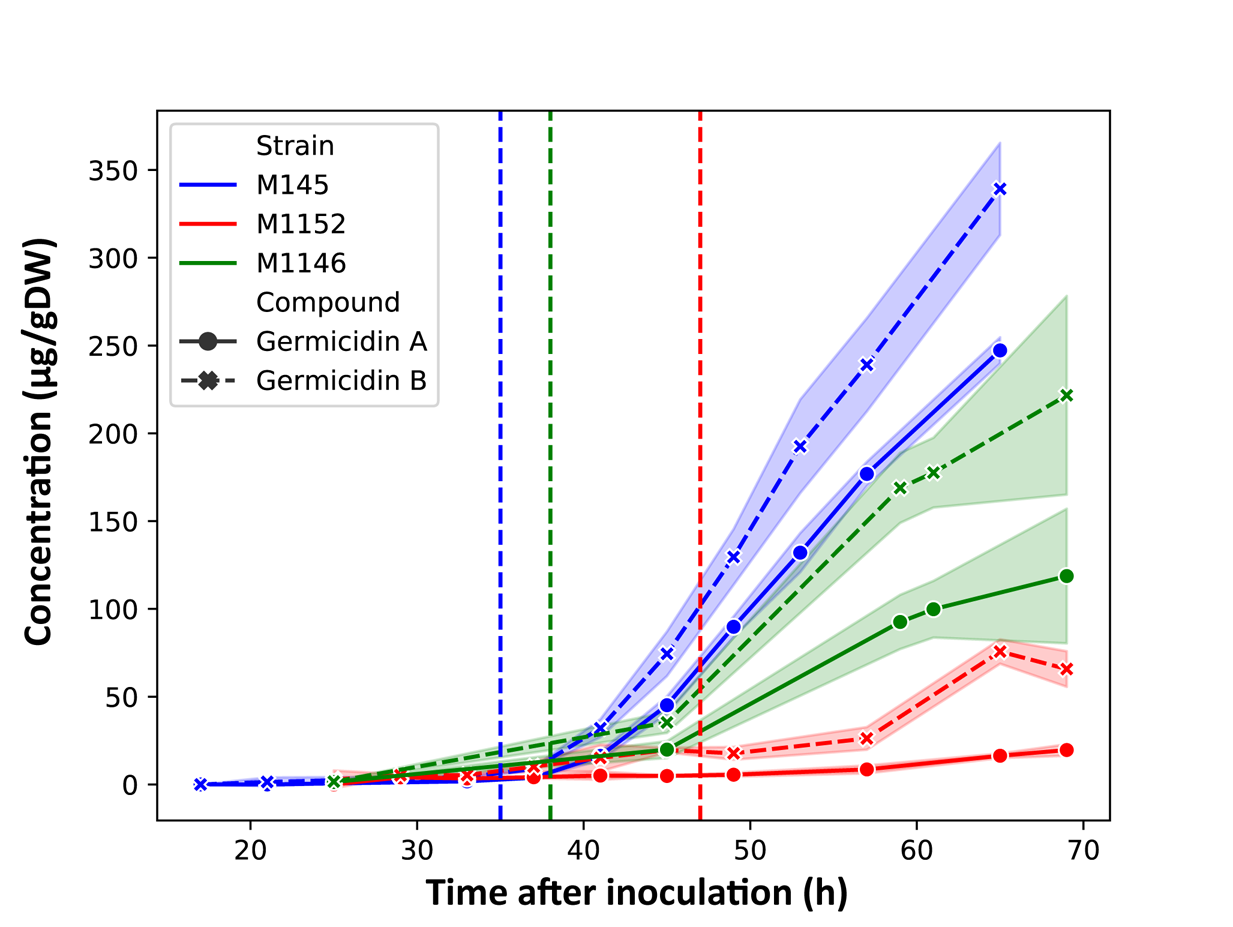
Concentrations of germicidin A and B produced by M145, M1146 and M1152. The concentrations are normalized by the biomass of each strain. The shaded regions display the uncertainty range (± 1 standard deviation) based on three replicate cultivations. Note that the growth rate is different between the strains, displayed by the vertical lines representing phosphate depletion at 35, 38 and 47 hours for M145, M1146 and M1152, respectively.

To further elucidate the cause of the reduced production in M1152, we also measured germicidin production in the intermediate strain M1146 (**Figure 7, Figure S7E**), which does not feature the *rpoB* mutation but is missing the 4 BGCs also deleted in M1152 (Gomez-Escribano and Bibb 2011). The production rate of germicidin A and B in M1146 was found to be reduced by 27% and 25%, respectively, compared to M145. In comparison to the strong reduction in germicidin production that can be assigned to the *rpoB* mutation in M1152, removal of only the 4 BGCs in M1146 has a moderate effect on germicidin production. This conforms with the minor contribution of the BGCs compared to fatty acid biosynthesis on the total consumption of malonyl-CoA (**Figure 5**). Nonetheless, it remains contradictory that the removal of polyketide precursor drains negatively impacts the production of other polyketides.

## Discussion

In this work, we carried out a multi-omics study to compare the metabolic changes of *Streptomyces coelicolor* M145 and the BGC-deletion mutant M1152 during batch fermentation. The defined cultivation medium used in this work was chosen because it supports sufficient growth and a delayed, well-defined onset of secondary metabolism, necessary to study the metabolic switch (Wentzel, Bruheim, et al. 2012). We aimed at defining the metabolic features differing between the two strains, both during exponential growth and stationary phase after phosphate depletion.

To achieve this from a systems biology perspective, we combined time-course sampled cultivation and transcriptome analysis with enzyme-constrained genome-scale models generated with proteome data. Such genome-scale models are extensively used to connect transcriptome- and proteome data to metabolic fluxes. Leveraging metabolic simulations to contextualize transcriptional changes is mainly impacted by the quality of the computational model used. Here, two teams joined efforts to improve a consensus model of *S. coelicolor*, yielding a comprehensive model useful for the scientific community.

### Genome-scale models provide hypothesis for slow growth of M1152

The reduced growth rate of M1152 is correlated with reduced glucose uptake and enhanced glutamate uptake compared to M145. This is expected to lead to a less active glycolysis but a more active TCA cycle, and thus, a more active oxidative metabolism in M1152 compared to M145. An active oxidative metabolism is known to generate oxidative stress, and indeed, the *in vivo* data, as well as the genome-scale model, predict an increased oxidative stress in M1152. The toxicity of oxidative stress might, at least in part, be responsible for the growth delay of M1152, while the *rpoB* mutation may add to this phenotype, since one of the functions of the ppGpp-associated RNA polymerase is to promote a growth arrest in conditions of nutritional stress.

### Further development may improve M1152 as host for heterologous expression

The strain M1152 has several advantages as a host for heterologous production of secondary metabolites. The deletion of the 4 major BGCs not only removes presumed competing drains for valuable precursors, but also generates a clean background to ease the identification of novel products by mass spectrometry. M1152 was already proven to be more efficient than M145 and M1146 in heterologous production of the nitrogen-containing antibiotics chloramphenicol and congocidine, as well as Act production from reintroduction of its BGC (Gomez-Escribano and Bibb 2011). Strains M1146 and M1152 produce, respectively, 3- to 5-, and 20- to 40-fold more chloramphenicol and congocidine from respective heterologous clusters than M145, a clear demonstration of the huge impact on production due to the *rpoB* mutation. While this contrasts with our data showing that M1152 has the lowest production of germicidin, it is relevant to note that chloramphenicol and congocidine are non-ribosomal peptide synthases relying on amino acids rather than malonyl-CoA as precursors. Although our data shows reduced degradation of branched-chain amino acids and metabolism of alanine, asparate and glutamate as the clearest metabolic divergence upon phosphate depletion in M1152, since congocidine and chloramphenicol are based on aromatic amino acids the connection to increased production of these NRPs is not obvious. Another option is that the increased oxidative metabolism in M1152 provides more redox cofactors to drive the synthesis of these molecules. If competition for valuable precursors was rate-limiting, the absence of the polyketides actinorhodin and coelimycin P1 should at least enhance the production of germicidin, all being dependent on malonyl-CoA. Moreover, differences in cultivation media further convolute cross-study comparisons: the aforementioned study use a complex growth medium while we used a defined medium with glucose and glutamate, which has previously been optimized for studying the metabolic switch (Wentzel, Bruheim, et al. 2012).

Furthermore, (re-)introduction of a (secondary) copy of germicidin synthase gene *gcs* in strains M1152 and M1317—derived from M1152 by additional removal of three Type III PKS genes including *gcs*—gave a 7.8 and 10.7-fold increase in germicidin production, respectively, compared to M1152 with only the native copy of *gcs* (Thanapipatsiri et al. 2015). Thus, the largest increase in production is not achieved by removal of competing precursor drains, but rather effected by the re-introduction of *gcs*, probably because expression of the inserted gene is not constrained by the same regulatory mechanism as the native gene.

Although earlier work has suggested a competition for common precursors between fatty acids and secondary metabolites biosynthesis (Craney et al. 2012), our results suggest that other approaches than deletion of competing precursor drains may be more efficient in the development of an optimized expression host, and it seems likely that different classes of BGCs may require different hosts for maximal production. Our comparative analysis of M145 and M1152 supports this development, not only as a systemic description connecting non-trivial associations between phenotypic, genetic and metabolic differences, but also by highlighting cellular processes that seems to be out of balance in M1152. These includes upregulation of ribosomal genes, most likely an effect of the *rpoB* mutation, and increased oxidative metabolism and oxidative stress. Since Act itself works as an electron acceptor one may hypothesize that its presence could relieve some of this stress. Another approach is to reintroduce *scbR2* to avoid influencing the related regulators of development and secondary metabolism.

Although *S. coelicolor* seems to have a complex and not fully elucidated regulatory system several studies have shown that manipulation of regulatory genes can affect production of secondary metabolites (Jones et al. 2011; D.-J. Kim et al. 2003; Okamoto et al. 2003; Rodriguez et al. 2012). The complex regulation of secondary metabolite biosynthesis makes rational strain design difficult (Liu et al. 2013), but black-box approaches including random mutations and screening are still viable approaches for strain development (van den Berg et al. 2008; Crook and Alper 2012). The Sco-GEM can aid this development by predicting the impact of these genetic alterations and to interpret’omics data.

### Limitations of the Study

We have performed a thorough comparison, of *S. coelicolor* M1145 and M1152, but to fully attribute changes in metabolism to the different genetic modifications as well as to unravel possible epistatic interactions we believe that a comprehensive analysis that also include the intermediate strain M1146, and possibly also an M145 strain featuring only the *rpoB* mutation (Xu et al. 2002), will be necessary.

### Resource availability

#### Lead Contact

Further information and requests for resources and reagents should be directed to and will be fulfilled by the Lead Contact, Eduard J Kerkhoven.

#### Materials Availability

This study did not generate new unique reagents.

#### Data and Code Availability

The models and scripts generated during this study are available at GitHub (https://github.com/SysBioChalmers/Sco-GEM). Here, the latest version of the Sco-GEM is available in both YAML and SBML level 3 Version 1. Additionally, users can contribute to further model development by posting issues or suggest changes. The proteomics data have been deposited to the ProteomeXchange Consortium via the PRIDE (Perez-Riverol et al. 2019) partner repository with the dataset identifier PXD013178. Normalized proteome data is also available in **Data Set S1, Tab 14.** The transcriptomics data have been deposited in NCBI’s Gene Expression Omnibus and are accessible under accession number GSE132487 (M145) and GSE132488 (M1152). Normalized counts are also found in **Data Set S1, Tab 15.**

## Supporting information

Data Set 1

## Acknowledgements

The authors would like to acknowledge Bogdan I. Florea of Leiden University, Leiden, Netherlands, for running and monitoring the proteome measurements, and the bio-organic synthesis group at Leiden University for providing the opportunity to use their instrumentation. The authors would also like to acknowledge co-workers at SINTEF Industry, Trondheim, Norway: Ingemar Nærdal, Anna Lewin and Kari Hjelen for running the batch fermentations and Anna Nordborg, Janne Beate Øiaas and Tone Haugen for performing offline analyses and the germicidin analytics. The RNA-Seq sequencing was carried out by c.ATG, Tübingen, Germany.

This study was conducted in the frame of ERA-net for Applied Systems Biology (ERA-SysAPP) project SYSTERACT and the project INBioPharm of the Centre for Digital Live Norway (Research Council of Norway grant no. 248885), with additional support of SINTEF internal funding.

## Author contributions

Conceptualization, E.K., E.A., A.W., S.S., A.S.Y. and T.K. Methodology and software, E.K., S.S., A.S.Y. and T.K. Validation and formal analysis, C.D., K.N., D.V.D., S.S., T.K., E.K. Investigation, T.K., A.W., S.S., E.K. Data curation, S.S., T.K. Writing - original draft, S.S., T.K., D.V.D., C.D., K.N. Writing – review & editing, all authors. Visualization, S.S., E.K., C.D., D.V.D. Supervision, A.W., E.A., E.K., G.W. Project administration, A.W. Funding acquisition: A.W., E.K., E.A., G.W.

## Supplemental Figures

**Figure S1: Reaction subsystems and origin.** The number of reactions in Sco-GEM in each of the 15 subsystems, and from which model they originate from. The other reactions (orange) are added during reconstruction of Sco-GEM.

**Figure S2: Gene clusters associated with metabolic switch.** RNA-seq (left column) and proteomics (right column) from M145 of the 8 gene clusters associated with the metabolic switch as previously identified (Nieselt et al. 2010). The 8 clusters are: A) genes related to ribosomal proteins; B) genes related to nitrogen metabolism; C) Cpk gene cluster; D) genes related to development; E) genes upregulated in response to phosphate depletion; F) genes involved in synthesis of phosphate-free polymers; G) Act gene cluster; H) Red gene cluster

**Figure S3: Log-transformed expression levels of genes related to nitrogen metabolism.** The order of the genes is determined by hierarchical clustering to align genes with similar expression profiles next to each other. From the log2-transformed RNA-seq data we observe that glutamate import (SCO5774-5777), the glutamate sensing system *gluR-gluK* (SCO5778 and SCO57779), *glnR* (SCO4159) and *glnA* (SCO2198) are downregulated subsequent to phosphate depletion. The phosphate depletion occurs between the third and fourth time point, i.e. at 35 and 47 hours for M145 and M1152, respectively. We also observe that the first time point in M145 is very different from all other samples.

**Figure S4: Clustered heatmaps of Z-score based on CO2-normalized sum of fluxes of all pathways standardized within each pathway and separated into different subsystems / parts of the metabolism.** A) Central carbon metabolism. B) Amino acid metabolism. C) Metabolism of vitamins and cofactors. D) Pathways of Biosynthetic gene clusters. E) Lipid metabolism. F) Oxidative stress. G) Degradation of toxic compounds. H) All other pathways. For all panels only pathway with a minimum flux of 1e-8 mmol (g DW)^-1^ h^-1^ were included.

**Figure S5: RNA-seq, proteome and flux prediction of specific gene clusters and reactions.** A) This panel display increasing CO2-normalized flux through citrate synthase (CS) and isocitrate dehydrogenase (ICDHyr) at later time points in M145 as predicted by EcSco-GEM. These two reactions are both part of the TCA cycle, converting acetyl-CoA to citrate (citrate synthase) and isocitrate to alpha-ketogluterate (isocitrate dehydrogenase). B) Log2 normalized expression data of genes involved in oxidative phosphorylation for M145 (blue) and M1152 (orange). The average expression level is higher in M1152 than in M145 but increasing at later time points for both strains. The expression profiles are only partially overlapping along the x-axis (hours after inoculation) because of the reduced growth and therefore delayed cultivation of M1152. C-H) Comparison of log2 normalized expression data as calculated with (log2 M145)-log2(M1152), where positive values indicate upregulation in M145 relative to M1152, and vice versa for negative values. C) Increased expression of genes of the AfsR regulon in M145, while no significant difference in expression is observed for (D) ScbR regulon; (E) AdpA regulon; (F) ArgR regulon; (G) genes induced by ppGpp; and (H) genes repressed by ppGpp.

**Figure S6: Biomass yield on glucose and glutamic acid.** M1152 (orange) has a higher growth yield on glucose than M145 (blue). The yield on glutamic acid (dashed line) is similar between the two strains.

**Figure S7: Analysis of transcriptome data of genes.** A) The heatmap display the mean standardized log2 expression levels for the 7 clusters of differentially expressed genes as determined by unsupervised clustering (k-means). Cluster 1-3 are upregulated in M1152, while the last four (cluster 4-7) are upregulated from the beginning or at later time points in M145. B-D) Comparison of log2 normalized expression data as calculated with (log2 M145)- log2(M1152), where positive values indicate upregulation in M145 relative to M1152, and vice versa for negative values. B) Genes in the SoxR regulon are reducing expression in M1152 at later time points. C) Almost all genes in the ATP-synthase cluster are up-regulated in M1152 after the first time point. D) The transcription of ribosomal protein genes after the metabolic switch is increased in M1152 compared to M145. E) Batch cultivation data of *S. coelicolor* M1146, showing volume corrected respiration (CO2), phosphate (PO4) and cell dry weight (CDW). Error bars are standard deviations of three biological replicates.

**Figure S8: Gene Ontology enrichment analysis of the 7 clusters identified in the 499 differentially expressed genes, categorized by function into four clustered heatmaps**. Each heatmap shows the p-value for the enrichment of each GO-process. A) Genes related to reactive oxygen species, the ribosome or development process and cell wall formation. B) Oxireductase and iron / metal ion homeostasis. C) Regulation, biosynthesis and metabolism related to RNA and DNA. D) All other GO-annotations. E) This color palette is the legend for the column colors on top of each heatmap which displays which of the seven clusters each gene belongs to. The red palette covers cluster 1-3 (upregulated in M1152), while the blue palette covers cluster 4-7 (upregulated in M145). Note that no GO-processes were enriched for the genes in cluster 2.

**Figure S9: Comparison of normalization methods of randomly sampled fluxes**. Heatmap showing mean flux values normalized by A) total carbon uptake from glucose and glutamate, B) CO2 production, C) sum of all fluxes and D) growth rate. Because the mean flux values in these reactions are different by several orders of magnitude, we display the data as standardized values (for each reaction).

## Other Supplemental material

**Supplemental Information:** The memote report of Sco-GEM and the Transparent Methods are provided in the Supplemental Information.

## Data Set S1

**Tab 1:** Detailed overview of the script performing the reconstruction of Sco-GEM.

**Tab 2:** Comparison of the new biomass reaction in Sco-GEM with the biomass reaction in iAA1259.

**Tab 3:** Reversibility prior update, calculated change in Gibbs free energy and standard deviation of the calculated change in Gibbs free energy of 770 reactions in Sco-GEM.

**Tab 4:** Overview of all transport reactions added or curated during the process of Sco-GEM model development, and the metabolites added along with the new transport reactions.

**Tab 5:** Genes present in the 7 clusters identified with K-Means clustering of the differentially expressed genes.

**Tab 6:** Sco4 ID, name and Sco-GEM ID of the reactions added from Sco4 to Sco-GEM.

**Tab 7:** Sco4 ID, name and Sco-GEM ID of the metabolites added from Sco4 to Sco-GEM

**Tab 8:** Reaction ID, name and gene annotation of reactions added from iAA1259 to Sco- GEM.

**Tab 9:** List of reactions modified according to iAA1259.

**Tab 10**: Metabolite ID and name of metabolites added from Sco4 to Sco-GEM.

**Tab 11**: List of SBO terms used in Sco-GEM

**Tab 12**: Single, pair and triplets of reactions which reduced model accuracy, in total 56 different reactions. The reversibility of these reactions was not changed according to the predicted change in Gibbs free energy.

**Tab 13**: Manual curation of the reversibility of the 82 reactions involving ATP predicted to be reversible by eQuilibrator.

**Tab 14**: Normalized proteome data for M145 and M1152. M145 is cultivated in F516-F518, while M1152 is cultivated in F519, F521, F522. Re-index IDs corresponds to the strain, fermentation-number (F516-F522), a unique number describing the experiment, and a numbering of the time points from 1 to 9 (P1-P9). I.e. the Re-indexed ID M145_F516_P2 represents the second timepoint of fermentation 516 with the strain M145. In the original IDs (row below) the P-number represents time after inoculation (in hours).

**Tab 15**: Normalized RNA-seq data for all six fermenters. M145 is cultivated in F516-F518, while M1152 is cultivated in F519, F521, F522. Re-index IDs corresponds to the strain, fermentation-number (F516-F522), a unique number describing the experiment, and a numbering of the time points from 1 to 9 (P1-P9). I.e. the Re-indexed ID M145_F516_P2 represents the second timepoint of fermentation 516 with the strain M145. In the original IDs (row below) the P-number represents time after inoculation (in hours).

**Tab 16**: List of genes within each cluster known to be associated with the metabolic switch.

