## Supplemental information for "Enzyme-constrained models and omics analysis of Streptomyces coelicolor reveal metabolic changes that enhance heterologous production"

The Supplemental Information contains four sections:

1. memote snapshot report of Sco-GEM
2. Transparent Methods

#### memote snapshot report of Sco-GEM

The following report (next page) was prepared by running memote version 0.9.12 in the command-line from the directory of the cloned Sco-GEM repository with the command: *memote report snapshot –custom-tests ComplementaryScripts/tests*

### Independent Section

Contains tests that are independent of the class of modeled organism, a model's complexity or types of identifiers that are used to describe its components. Parameterization or initialization of the network is not required. See readme for more details.

#### Consistency

| | |
|---|---|
| Stoichiometric Consistency | 37.9% |
| Mass Balance | 84.6% |
| Charge Balance | 96.8% |
| Metabolite Connectivity | 100.0% |
| Unbounded Flux In Default Medium | 84.9% |
| Sub Total | 69% |

#### Annotation - Metabolites

| | |
|---|---|
| Presence of Metabolite Annotation | 100.0% |
| Metabolite Annotations Per Database | Info |
| pubchem.compound | 18.5% |
| kegg.compound | 82.4% |
| seed.compound | 1.6% |
| inchikey | 0.0% |
| inchi | 18.6% |
| chebi | 80.3% |
| hmdb | 0.0% |
| reactome | 0.0% |
| metanetx.chemical | 88.2% |
| bigg.metabolite | 95.9% |
| biocyc | 70.2% |

### Specific Section

Covers general statistics and specific aspects of a metabolic network that are not universally applicable. See readme for more details.

#### SBML

| | |
|---|---|
| SBML Level and Version | SBML Level 3 Version 1 |
| FBC enabled | true |

#### Basic Information

| | |
|---|---|
| Model Identifier | |
| Total Metabolites | 2,073 |
| Total Reactions | 2,612 |
| Total Genes | 1,778 |
| Total Compartments | 2 |
| Metabolic Coverage | 1.47 |

#### Metabolite Information

| | |
|---|---|
| Unique Metabolites | 1,836 |
| Duplicate Metabolites in Identical Compartments | 0 |
| Metabolites without Charge | 0 |
| Metabolites without Formula | 0 |
| Medium Components | 22 |

#### Reaction Information

| | |
|---|---|
| Purely Metabolic Reactions | 2,011 |
| Purely Metabolic Reactions with Constraints | 1 |
| Transport Reactions | 325 |

| | |
|---|---|
| kegg.compound | 100.0% |
| seed.compound | 100.0% |
| inchikey | 0.0% |
| inchi | 100.0% |
| chebi | 99.9% |
| hmdb | 0.0% |
| reactome | 0.0% |
| metanetx.chemical | 99.8% |
| bigg.metabolite | 100.0% |
| biocyc | 99.4% |
| Uniform Metabolite Identifier Namespace | 100.0% |
| Sub Total | 79% |

#### Annotation - Reactions

| | |
|---|---|
| Presence of Reaction Annotation | 100.0% |
| Reaction Annotations Per Database | Info |
| rhea | 37.4% |
| kegg.reaction | 52.4% |
| seed.reaction | 0.0% |
| metanetx.reaction | 79.2% |
| bigg.reaction | 93.6% |
| reactome | 0.0% |
| ec-code | 64.4% |
| brenda | 0.0% |
| biocyc | 44.3% |
| Reaction Annotation Conformity Per Database | Info |

| | |
|---|---|
| Reactions With Partially Identical Annotations | 0.09 |
| Duplicate Reactions | 0.00 |
| Reactions With Identical Genes | 0.48 |

#### Gene-Protein-Reaction (GPR) Associations

| | |
|---|---|
| Reactions without GPR | 342 |
| Fraction of Transport Reactions without GPR | 0.26 |
| Enzyme Complexes | 199 |

#### Biomass

| | |
|---|---|
| Biomass Reactions Identified | 10 |
| Biomass Consistency | Info |
| MISC_PSEUDO | Errored |
| CARBOHYDRATE_PSEUDO | Errored |
| PROTEIN_PSEUDO_tRNA | Errored |
| LIPID_PSEUDO | Errored |
| CELL_WALL_PSEUDO | Errored |
| BIOMASS_SCO_tRNA | Errored |
| DNA_PSEUDO | Errored |
| RNA_PSEUDO | Errored |
| BIOMASS_SCO | Errored |
| PROTEIN_PSEUDO | Errored |
| Biomass Production In Default Medium | Info |
| MISC_PSEUDO | 0.07 |
| CARBOHYDRATE_PSEUDO | 0.07 |
| PROTEIN_PSEUDO_tRNA | 0.07 |
| LIPID_PSEUDO | 0.07 |

| | |
|---|---|
| seed.reaction | 0.0% |
| metanetx.reaction | 100.0% |
| bigg.reaction | 100.0% |
| reactome | 100.0% |
| ec-code | 97.2% |
| brenda | 0.0% |
| biocyc | 100.0% |
| Uniform Reaction Identifier Namespace | 100.0% |
| Sub Total | 80% |

#### Annotation - Genes

| | |
|---|---|
| Presence of Gene Annotation | 89.5% |
| Gene Annotations Per Database | Info |
| refseq | 89.5% |
| uniprot | 89.5% |
| ecogene | 0.0% |
| kegg.genes | 0.0% |
| ncbigi | 0.0% |
| ncbigene | 0.0% |
| ncbiprotein | 0.0% |
| ccds | 0.0% |
| hprd | 0.0% |
| asap | 0.0% |
| Gene Annotation Conformity Per Database | Info |
| refseq | 11.1% |
| uniprot | 100.0% |

| | |
|---|---|
| DNA_PSEUDO | 0.07 |
| RNA_PSEUDO | 0.07 |
| BIOMASS_SCO | 0.07 |
| PROTEIN_PSEUDO | 0.07 |
| Unrealistic Growth Rate In Default Medium | Info |
| MISC_PSEUDO | false |
| CARBOHYDRATE_PSEUDO | false |
| PROTEIN_PSEUDO_tRNA | false |
| LIPID_PSEUDO | false |
| CELL_WALL_PSEUDO | false |
| BIOMASS_SCO_tRNA | false |
| DNA_PSEUDO | false |
| RNA_PSEUDO | false |
| BIOMASS_SCO | false |
| PROTEIN_PSEUDO | false |
| Biomass Production In Complete Medium | Info |
| MISC_PSEUDO | 123.45 |
| CARBOHYDRATE_PSEUDO | 123.45 |
| PROTEIN_PSEUDO_tRNA | 123.45 |
| LIPID_PSEUDO | 123.45 |
| CELL_WALL_PSEUDO | 123.45 |
| BIOMASS_SCO_tRNA | 123.45 |
| DNA_PSEUDO | 123.45 |
| RNA_PSEUDO | 123.45 |
| BIOMASS_SCO | 123.45 |
| PROTEIN_PSEUDO | 123.45 |

| | |
|---|---|
| ncbigi | 0.0% |
| ncbigene | 0.0% |
| ncbiprotein | 0.0% |
| ccds | 0.0% |
| hprd | 0.0% |
| asap | 0.0% |
| Sub Total | 40% |

#### Annotation - SBO Terms

| | |
|---|---|
| Metabolite General SBO Presence | 100.0% |
| Metabolite SBO:0000247 Presence | 99.6% |
| Reaction General SBO Presence | 100.0% |
| Metabolic Reaction SBO:0000176 Presence | 99.8% |
| Transport Reaction SBO:0000185 Presence | 100.0% |
| Exchange Reaction SBO:0000627 Presence | 100.0% |
| Demand Reaction SBO:0000628 Presence | 100.0% |
| Sink Reactions SBO:0000632 Presence | Skipped |
| Gene General SBO Presence | 89.5% |
| Gene SBO:0000243 Presence | 89.5% |
| Biomass Reactions SBO:0000629 Presence | 100.0% |
| Sub Total | 89% |

| | |
|---|---|
| Total Score | 77% |

Total Score

| | |
|---|---|
| CARBOHYDRATE_PSEUDO | 0 |
| PROTEIN_PSEUDO_tRNA | 20 |
| LIPID_PSEUDO | 0 |
| CELL_WALL_PSEUDO | 0 |
| BIOMASS_SCO_tRNA | 0 |
| DNA_PSEUDO | 0 |
| RNA_PSEUDO | 0 |
| BIOMASS_SCO | 0 |
| PROTEIN_PSEUDO | 0 |
| Blocked Biomass Precursors In Complete Medium | Info |
| MISC_PSEUDO | 0 |
| CARBOHYDRATE_PSEUDO | 0 |
| PROTEIN_PSEUDO_tRNA | 20 |
| LIPID_PSEUDO | 0 |
| CELL_WALL_PSEUDO | 0 |
| BIOMASS_SCO_tRNA | 0 |
| DNA_PSEUDO | 0 |
| RNA_PSEUDO | 0 |
| BIOMASS_SCO | 0 |
| PROTEIN_PSEUDO | 0 |
| Ratio of Direct Metabolites in Biomass Reaction | Info |
| MISC_PSEUDO | 0.10 |
| CARBOHYDRATE_PSEUDO | 0.00 |
| PROTEIN_PSEUDO_tRNA | 0.00 |
| LIPID_PSEUDO | 0.00 |
| CELL_WALL_PSEUDO | 0.00 |

77%

Score per Category

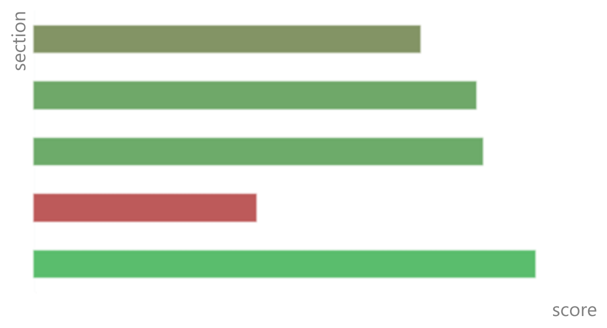


| | |
|---|---|
| RNA_PSEUDO | 0.00 |
| BIOMASS_SCO | 0.00 |
| PROTEIN_PSEUDO | 0.00 |
| Number of Missing Essential Biomass Precursors | Info |
| MISC_PSEUDO | 29 |
| CARBOHYDRATE_PSEUDO | 36 |
| PROTEIN_PSEUDO_tRNA | 37 |
| LIPID_PSEUDO | 37 |
| CELL_WALL_PSEUDO | 1 |
| BIOMASS_SCO_tRNA | 1 |
| DNA_PSEUDO | 1 |
| RNA_PSEUDO | 1 |
| BIOMASS_SCO | 1 |
| PROTEIN_PSEUDO | 17 |

#### Energy Metabolism

| | |
|---|---|
| Non-Growth Associated Maintenance Reaction | Errored |
| Growth-associated Maintenance in Biomass Reaction | Info |
| MISC_PSEUDO | false |
| CARBOHYDRATE_PSEUDO | false |
| PROTEIN_PSEUDO_tRNA | false |
| LIPID_PSEUDO | false |
| CELL_WALL_PSEUDO | false |
| BIOMASS_SCO_tRNA | false |
| DNA_PSEUDO | false |
| RNA_PSEUDO | false |
| BIOMASS_SCO | false |

| Erroneous Energy-generating Cycles | Info |
|---|---|
| MNXM3 | Skipped |
| MNXM63 | Skipped |
| MNXM51 | Skipped |
| MNXM121 | Skipped |
| MNXM423 | Skipped |
| MNXM6 | Skipped |
| MNXM10 | Skipped |
| MNXM38 | Skipped |
| MNXM208 | Skipped |
| MNXM191 | Skipped |
| MNXM223 | Skipped |
| MNXM7517 | Skipped |
| MNXM12233 | Skipped |
| MNXM558 | Skipped |
| MNXM21 | Skipped |
| MNXM89557 | Skipped |

#### Network Topology

| | |
|---|---|
| Universally Blocked Reactions | 683 |
| Orphan Metabolites | 140 |
| Dead-end Metabolites | 232 |
| Stoichiometrically Balanced Cycles | 156 |
| Metabolite Production In Complete Medium | 714 |
| Metabolite Consumption In Complete Medium | 876 |

#### Matrix Conditioning

| | |
|---|---|
| Rank | 1915 |
| Degrees Of Freedom | 697 |

#### Experimental Data Comparison

| | |
|---|---|
| Growth Prediction | Skipped |
| Gene Essentiality Prediction | Skipped |

#### Misc. Tests

| | |
|---|---|
| Test if all metabolites have been given a name | 1.00 |
| Test CDA production | 0.07 |
| Test germicidinB production | 0.37 |
| Test growth for knockout-mutants from the transposon mutagenesis study by Xu et al.(2017) | 0.76 |
| Test germicidinA production | 0.37 |
| Test RED production | 0.13 |
| Test germicidinC production | 0.33 |
| Test that the growth rate is around 0.075 | 0.07 |
| Test growth for knockout-mutants from the litterature in given environments | 0.63 |
| Test growth for WT in given environments | 0.96 |
| Test if all reactions have been given a name | 1.00 |
| Test ACT production | Errored |

#### Environment

| | |
|---|---|
| Python Version | 3.7.3 |
| Platform | Windows |
| Memote Version | 0.9.12 |

### Transparent methods

#### Sco-GEM consensus model reconstruction and development

Sco-GEM, the community consensus model for *Streptomyces coelicolor* is developed, maintained, hosted and publicly available on GitHub (https://github.com/SysBioChalmers/Sco-GEM). When we refer to files in the following sections, we use the file names and relative to the main folder in this GitHub repository. By hosting the model on GitHub, we make the reconstruction transparent, the data accessible, provide a structure framework for further development by the community. To this end we also created a channel on Gitter dedicated to Sco-GEM questions and discussions (https://gitter.im/SysBioChalmers/Sco-GEM). The model repository was created using memote (Lieven et al., 2018) and we use a GitFlow structure with two main branches, the *devel* branch contains the most recent changes while the *master* branch contains the stable releases. All new features or bug fixes are performed in separate branches that are incorporated into the *devel* branch through *pull requests.* Semantics for branch names and commit messages are described in *CONTRIBUTING.rst*. The main script language for the model reconstruction is python (version > 3.6), with the exception being the *feat/ecModel* branch with the development of the enzyme-constrained model (EcSco-GEM) where Matlab (version > 7.3) is used.

In terms of folder structure data files, scripts and model files are stored in *ComplementaryData*, *ComplementaryScripts*, and *ModelFiles*, respectively. In the main folder we find the following files:

- *.gitignore*: File which describes file formats automatically ignored by git
- *.gitconfig*: Git config file
- *.gitmodules*: List of linked submodules
- *CONTRIBUTING.rst*: Guidelines describing how to contribute
- *README.md*: General information about the repository
- *HISTORY.rst*: History of model version releases
- *LICENSE.md*: License information
- *memote.ini*: File created by memote (Lieven et al., 2018)
- *requirements.txt*: List of python-packages required to run the model reconstruction

- *.travis.yml*: Config file for automatization of memote with Travis (https://travis-ci.org/)

Sco-GEM can be reconstructed at any time using the python script *ComplementaryScripts/reconstruct_scoGEM.py*. Each task of the reconstruction process is performed in a separate script and associated with an issue on GitHub (**Data Set S1**, **Tab 1**). The details of each task are described in the following paragraphs.

##### Curate identified issues in iKS1317

We used iKS1317 (Kumelj et al., 2018) as the starting point for the reconstruction of Sco-GEM. Since the publication of iKS1317, several issues had been identified and these were curated as the initial step in the reconstruction pipeline. The curations include correcting the mass and charge balance of the reactions NOR_syn, OAADC, SEPHCHCS and DIOP5OR, and correcting the ec-code, KEGG annotation and gene association for the reactions 3OXCOAT, MMSYNB, PGMT, PPM, ME1, GLUDyi, GLUSx, GLUSy and GLUN.

##### Curate and add reactions from Sco4

The Sco4 GEM of *S. coelicolor* (Wang et al., 2018) contained additional reactions that we wanted to include in Sco-GEM. However, prior to adding content from Sco4 we curated issues that had been identified since publication. Eleven reactions were found to be either duplicated or wrong in Sco4, and these were removed: RXN0-5224, METHYLGLUTACONYL-COA-HYDRATASE-RXN, GLU6PDEHYDROG-RXN, RXN-15856, 1.14.13.84-RXN_NADPH, R03998, R03999, R09692_NADPH, RXN-9930, 1.17.1.1-RXN_NADH, R09692_NADH. We additionally updated the gene annotations of the following reactions: RMPA, ABTDG, PROD2, THRPDC, ADCL, OXPTNDH, GLNTRS, CU2abc, CBIabc, CBL1abc, GSNt2, INSt2 and PDH.

To enable addition of reactions from Sco4 (Wang et al., 2018) to Sco-GEM we mapped reactions added during the Sco4 development to reactions present in iKS1317 (Kumelj et al., 2018). This mapping was performed semi-automatically: automatic mapping using KEGG and BioCyc annotations followed by manual curation. In total, 394 new reactions and 404 new metabolites were added from Sco4 to Sco-GEM (**Data Set S1, Tab 6 and 7**). Most of the reactions and metabolites added from Sco4 had IDs from the MetaCyc database (Caspi et al., 2014), containing characters such as dash or parentheses not properly handled by the SBML parser in COBRApy (Ebrahim et al., 2013). Thus, the ID of all reactions and metabolites added from Sco4 were changed to the correct BiGG ID if possible, otherwise a new ID was created according to the guidelines given in BiGG (King et al., 2016). KEGG (Kanehisa, 2000) and MetaNetX (Moretti et al., 2016) identifiers were included as annotations when possible. Full lists of the IDs and annotations given to reactions and metabolites added from Sco4 are found in the GitHub repository folder *ComplementaryData/curation* as *added_sco4_reactions.csv* and *added_sco4_metabolites.csv*, respectively.

##### Add gene annotations, reactions and metabolites to Sco-GEM from iAA1259

Based on supplementary files 4 and 5 from iAA1259 (Amara et al., 2018) which list the reactions and metabolites added in iAA1259, we identified 44 reactions and 31 metabolites present in neither Sco4 or iKS1317 (**Data Set S1, Tab 8 and 10**). These 44 reactions were added from iAA1259 and were mainly related to coelimycin biosynthesis, xylan and cellulose degradation and butyrolactones pathway. We further incorporated the modification of 27 reactions curated in iAA1259, associated with oxidative phosphorylation, futalosine pathway or chitin degradation (**Data Set S1, Tab 9**). These curations mainly updated gene-reaction rules but also updated reaction bounds and deletion of two reactions (CFL and DHFUTALS). Finally, we incorporated the biomass-function which was updated in iAA1259.

##### Change direction of reactions that were backwards irreversible

The pipeline for reconstruction of the enzyme-constrained model required all reactions to be either reversible or forward irreversible (i.e. reactions with bounds (-1000, 0) are not allowed). Therefore, all backward irreversible reactions were rewritten (substrates were changed to products and *vice versa*) so they could be represented as forward irreversible.

##### Fix missing / wrongly annotated reactions and metabolites

We identified several minor issues related to reaction and metabolite IDs or annotations. These may come from the current or previous model reconstruction efforts. These issues include:

- Misspelled IDs or annotations

- Empty annotations in SBML file
- Wrong BioCyc annotations for metabolites and reactions in the germicidin pathway
- Update all MetaNetX annotations
- Exchange reactions given BioCyc annotations
- Fix chebi annotations so they comply with the MIRIAM identifiers
- Mixed up IDs for actACPmmy and malACPmmy

##### Create pseudo-metabolites for NADH/NADPH and NAD+/NADP+ to use in reaction were the redox cofactor is not known

For some redox reactions added from Sco4, it was not sure if NADH/NAD+ or NADPH/NADP+ was the participating cofactor pair. In this case, both possibilities were included in Sco4. However, to avoid duplicated reactions and make it explicit that the cofactor is unknown we changed these reactions to use pseudo-metabolites (acceptor_c and donor_c) as the cofactor pair. We then also included pseudo-reactions which converts NADH/NADPH and NAD+/NADP+ to donor_c and acceptor_c, respectively [pseudo-reaction IDs: PSEUDO_DONOR_NADH; PSEUDO_DONOR_NADPH; PSEUDO_ACCEPTOR_NAD; PSEUDO_ACCEPTOR_NADP). In total 17 enzymatic reactions use these pseudo-metabolites as cofactor pair: 3OCHOCDH; OXCOADH; 4DPCDH; 4HYDPRO; 4NITROB; AHLGAL; AHOPS; CADHX; DDALLO; DPCOX; GDP64HRD; HDAPMO; PHYFLUDS; HYTDES; SORBDH; ZCARDS; ZCAROTDH2.

##### Add SBO terms to genes, reactions and metabolites

SBO (Systems Biology Ontology) (Courtot et al., 2011) terms were included as annotations of reactions, genes and metabolites according to **Data Set S1, Tab 11**.

##### Update the biomass reaction

In iAA1259, the biomass reaction was curated in respect to 2-demethylmenaquinol and menaquinol, however, this resulted in a biomass reaction that combined described more than 1 g per gDCW. In addition, the biomass reaction of all *S. coelicolor* models have described small molecule and protein co-factors/prosthetic groups as components, where their abundance was arbitrarily set to complement the remaining biomass components to reach 1 g per gDCW. This is likely a gross overestimation for many of these molecules, and this proved problematic for initial simulations with the enzyme constrained model. In contrast to enzymes of central carbon metabolism, enzymes involved in biosynthesis of such co-factors and prosthetic groups have typically lower efficiency, such that large fractions of the protein allocation would have to be devoted to these pathways if the abundances are overestimated.

The availability of proteomics data has allowed us to give more reasonable estimates of abundance of protein-linked cofactors and prosthetic groups. The new biomass reaction was estimated through the following steps:

1. By querying UniProt, a list of prosthetic groups per protein were collated (*ComplementaryData/biomass/prosthetic_groups_uniProt.txt*) and further processed (*ComplementaryScripts/ecModel/prostheticGroups.m*) as detailed below.
2. If *metal* was specified as cofactor, the abundance was split over cobalt$^{2+}$, cupper$^{2+}$, iron$^{2+}$, zinc$^{2+}$, nickel$^{2+}$, calcium$^{2+}$, potassium$^{+}$, magnesium$^{2+}$ and manganese$^{2+}$.
3. Dipyrromethane is generated by the enzyme itself from its substrate and is therefore not further considered.
4. From the M145 and M1152 cultivation data, quantitative proteomics was estimated as detailed below.
5. Cofactor abundances were estimated by combining the estimated protein levels and the protein cofactor annotation (available at *ComplementaryData/biomass/ prosthetic_groups_mets.txt*)
6. To simplify fitting of biomass components, the full biomass reaction was split into the pseudometabolites *lipid*, *dna*, *rna*, *protein*, *carbohydrate, cell_wall* and *misc*, with the latter containing the cofactors (*ComplementaryData/biomass/standard_biomass.txt*).
7. After updating the abundances of the cofactors, the remaining *misc* metabolites were $_r$refitted to ensure that the total biomass adds up to 1 g per gDCW.

The updated composition (*ComplementaryData/biomass/biomass_scaled.txt*) was subsequently used to modify the model stoichiometry (*fix_biomass.py*). A comparison of the updated biomass reaction and the biomass reaction in iAA1259 is presented in **Data Set S1, Tab 2**.

#### Model reversibility

By using the python-API (https://gitlab.com/elad.noor/equilibrator-api) of eQuilibrator (Flamholz et al., 2012) we calculated the change in Gibbs free energy for 770 reactions (**Data Set S1, Tab 3**). eQuilibrator can only calculate the change in Gibbs free energy for intracellular reactions (i.e. not transport and exchange reactions) where all metabolites are mapped to KEGG (Kanehisa, 2000; Kanehisa et al., 2019). The calculations are based on the component contribution method (Noor et al., 2013). The change in Gibbs free energy was calculated at standard conditions (25 °C, 1 bar), pH7 and 1mM concentration of reactants, denoted $\Delta_r G'^m$ in eQuilibrator. We then applied a threshold of -30 kJ/mol to define a reaction as irreversible (Bar-Even et al., 2012; Feist et al., 2007), and compared the calculated reversibility with the reversibility of these reactions in the model prior to curation. We found that the reversibility was equal for 56.9% (438 / 770) of the reactions (**Figure 1E**). The majority of differences were reactions that were irreversible in the model but classified as reversible using the calculated values for the change in Gibbs free energy (35%; 273/770; **Figure 1E**).

Using the set of growth data and knockout data, we evaluated the effect of the suggested changes in reaction reversibility: by randomly applying these changes to 10 reactions at the time, we identified 13 single, 22 pairs and 13 triplets of reactions (consisting of 55 unique reactions) that reduced model accuracy when the reversibility was changed based on the change in Gibbs free energy (**Data Set S1, Tab 12**). Then we used the data set of growth and gene knockout phenotypes (Kumelj et al., 2018) to identify another 6 reactions that caused erroneous predictions if the reversibility were changed (PROD2, ARGSS, OCT, URIK1, URIK2, and UPPRT). These 61 reactions were discarded from having the reversibility changed, resulting in a total of 271 reactions with changed reversibility.

Energetic cofactors, including ATP, NADPH, NADH, FAD and any quinone, were involved in 284 of the 770 reactions for which the change in Gibbs free energy was calculated. Of the 114 reactions involving ATP, 82 reactions had an estimated change in Gibbs free energy between ±30 kJ/mol, indicating that the reactions were reversible. Because one assumes that ATP-driven reactions in general are irreversible (Thiele and Palsson, 2010), the reversibility of these 82 reactions were manually curated (**Data Set S1, Tab 13**). For the 7 quinone-associated reactions for which the change in Gibbs free energy was calculated (CYTBD2, NADH17b, NADH10b, MBCOA2, G3PD5, PDH3, NADH2r) all were defined as irreversible as previously suggested (Thiele and Palsson, 2010). The reversibility of reactions involving any of the other energetic cofactors were treated as any other reaction as previously described.

##### Analysis and annotation of transport reactions

Gene annotations, substrate and transport class information were mostly extracted from Transport DB 2.0 (Elbourne et al., 2017) and TCDB (Saier et al., 2016). Then, transport proteins were extracted from IUBMB-approved Transporter Classification (TC) System and categorized into 9 main classes (**Figure 1F**): 1) ABC transporter; 2) PTS transporter; 3) Proton symporter; 4) Sodium symporter; 5) Other symporter; 6) Proton antiport; 7) Other antiport; 8) Facilitated diffusion; 9) Simple diffusion. For those transport proteins with an ambiguous substrate annotation in TCDB, the specific substrate annotation was obtain by extracting annotations from KEGG (Kanehisa, 2000; Kanehisa et al., 2019), UniProt (The UniProt Consortium, 2019) or through BLAST homology search (NCBI Resource Coordinators, 2017) using a similarity threshold of 90% (**Data Set S1, Tab 4**).

##### Subsystem annotations

We leveraged the KEGG and BioCyc annotations of each individual reaction to extract a draft subsystem and pathway annotation for each reaction. For KEGG, this was achieved by using the python module BioServices (Cokelaer et al., 2013) while we used PythonCyc (https://github.com/latendre/PythonCyc) and PathwayTools (Karp et al., 2016) to extract pathway annotations from BioCyc (Karp et al., 2017).

The draft annotations were then curated, and each reaction was annotated to one out of 15 subsystems. When no or multiple annotations were extracted from the databases we used adjacent reactions in the metabolic network to infer the single, most correct annotation. These 15 subsystem categories are based on the categories of the KEGG Pathway Maps for metabolism (https://www.genome.jp/kegg/pathway.html) but we have included three additional categories to cover all aspects of the model: Biomass and maintenance functions, Membrane Transport, and Exchange (**Figure S1**).

We also annotated 1964 of the 2552 reactions to one out of 128 different pathways. The remaining 588 are mostly transport and exchange reactions, or possibly reactions not fitting into any of these pathways.

##### Export model file with alphabetical ordering

An import feature with GitHub is the ability to easily see changes in text files after every commit. However, COBRApy (Ebrahim et al., 2013) doesn’t sort the list of reactions, metabolites and genes before the SBML-file is written and this make it look like there was a lot of changes even when the model is unchanged. Thus, we now sort these lists before writing to file. The export function also stores the model-file in the YAML format which is more readable than SBML (XML). Finally, the export function creates the *requirements.txt* file which holds information about all non-standard python modules necessary to run the model reconstruction.

#### Development of enzymatically constrained (EcSco-GEM) model

An enzyme-constrained version of the Sco-GEM model (denoted EcSco-GEM) was generated using GECKO (Sánchez et al., 2017). The GECKO method enhances an existing GEM by explicitly constraining the maximum flux through each reaction by the maximum capacity of the corresponding enzyme, given by the product of the enzyme abundance and catalytic coefficient. Both reversible reactions and reactions catalysed by isoenzymes (redundant genes) are handled automatically by the GECKO method by splitting each occurrence into individual reactions. The Sco-GEM v1.1 model was modified using GECKO version 1.3.4. Kinetic data, in the form of $k_{cat}$ values ($s^{-1}$), were automatically collected from BRENDA (Jeske et al., 2019). If BRENDA did not report a $k_{cat}$ value for an enzyme, GECKO searched for alternative $k_{cat}$ values by reducing specificity, on the level of substrate, enzymatic activity (EC number) and organism.

A total of 4753 $k_{cat}$ values were matched, including separate values for forward and backward direction for reversible reactions, of which:

- 53 were matched with organism (S. coelicolor) and correct substrate
- 1541 were matched with closest organism and correct substrate
- 236 were matched with organism (S. coelicolor) and any substrate

- 15 were matched with organism (S. coelicolor) and any substrate, reported specific activities instead of $k_{cat}$ (corrected in the model for molecular weight of the enzyme)
- 2586 were matched with closest organism and any substrate
- 322 were matched with any organism and any substrate, reported specific activities instead of $k_{cat}$ (corrected in the model for molecular weight of the enzyme)

The algorithm first looped through these criteria above, with the full EC code. If no match could be found, wildcards were added (e.g. EC2.3.4.- instead of EC2.3.4.5), followed by going through the list of criteria above. The statistics there is:

- 4178 were matched without any wildcards (full EC code)
- 544 were matched after adding one wildcard
- 21 were matched after adding two wildcards (e.g. EC2.3.-.-)
- 10 were matched after adding three wildcards
- 0 were matched after adding four wildcards

Using the initial set of BRENDA-suggested $k_{cat}$ values, the model was evaluated to support simulation of experimentally measured growth rates. During this testing the NAD(H)/NAD(P)H pseudo-reactions were blocked to avoid infeasible loops.

The following 6 $k_{cat}$ values were identified as growth limiting resulting in the stated manual curations:

- <u>Chorismate synthase</u> (CHORS; EC4.2.3.5; SCO1496; Q9KXQ4)
  Sco-GEM uses 5-O-(1-Carboxyvinyl)-3-phosphoshikimate as name of the main substrate, while BRENDA uses its synonym 5-enolpyruvylshikimate 3-phosphate. This prevented automatically finding the substrate. Hence, the kcat was manually changed to 0.87 $s^{-1}$, as measured from *N. crassa* (Rauch et al., 2008).

- <u>Phosphoribosylformylglycinamidine synthase</u> (PRFGS; EC6.3.5.3; SCO4077 and SCO4078 and SCO4079; Q9RKK5 and Q9RKK6 and Q9RKK7)

$k_{cat}$ suggested by BRENDA used NH4+ as substrate, instead of glutamine. Specific activity using glutamine is provided for *E. coli*: 2.15 µmol/min/mg protein (Schendel et al., 1989). Assuming molecular weight of 141 kDa, this translates to $k_{cat}$ = 5.05 $s^{-1}$.

- Methylmalonate-semialdehyde dehydrogenase (malonic semialdehyde) (MMSAD3; EC1.2.1.27; SCO2726; Q9L1J1)
  $k_{cat}$ suggested by BRENDA is from archaea, instead use $k_{cat}$ value of 2.2 $s^{-1}$ from *B. subtilis* (Talfournier et al., 2011).

- Phosphoribosyl-ATP pyrophosphatase (PRATPP; EC3.6.1.31; SCO1439; Q9EWK0)
  $k_{cat}$ suggested by BRENDA was calculated from specific activity in *Salmonella enterica*, but the reported value was measured in cell extract, not from purified enzyme. Instead, use specific activity from *S. cerevisiae*: 332 umol/min/mg protein (Keesey et al., 1979). Assuming molecular weight of 95 kDa, this translates to a $k_{cat}$ of 526 $s^{-1}$.

- Glyceraldehyde 3-phosphate dehydrogenase (Q9Z518/EC1.2.1.12) - assigned $k_{cat}$ from Corynebacterium glutamicum was highly growth limiting. Instead use specific activity measured of pentalenolactone sensitive gapdh in Streptomyces arenae: 112 umol/min/mg protein (Maurer et al., 1983).

Then, separate models were created for each strain (the gene clusters for actinorhodin, undecylprodigiosin, CDA and coelimycin P1 were removed to create M1152) and for each time point by using estimated growth, uptake rates of glutamate and glucose, secretion rates of undecylprodigiosin, germicidin A and B and proteome measurements. The estimated growth, uptake and secretion rates were estimated from raw measurements across three biological replicates (details provided in the last section). These time point specific models (9 time points for M145, 8 time points for M1152) were used to analyse the activity in individual metabolic pathways through random sampling (Bordel et al., 2010). We also created one EcSco-GEM model for each strain with a global constraint on the protein usage instead of specific protein usage, which were used for model quality control.

#### Continuous integration and quality control with memote

Validation and quality assessment of Sco-GEM is carried out using the test-suite in memote (Lieven et al., 2018). Memote provides by default a large range of tests, which we have used to identify issues and possible improvements. The test suite reports descriptive model statistics such as the number of genes, reactions and metabolites, and also checks the presence of SBO terms and annotations, the charge and mass balance of all reactions, the network topology and find energy-generating cycles (Fritzemeier et al., 2017). Additionally, we incorporated custom tests into the memote test-suite to automatically compare predicted phenotypes with experimental data in different growth media and for different knockout mutants. In addition to the classical binary classifiers accuracy, sensitivity and specificity we also report the Matthews correlation coefficient which is considered to be more reliable when the number of elements in each classification category is skewed (Chicco and Jurman, 2020). The Matthews correlation coefficient (*MCC*) is calculated from the true positive (*TP*), false positive (*FP*), true negative (*TN*) and false negative (*FN*) values as $MCC = \frac{TP \cdot TN - FP \cdot FN}{\sqrt{(TP+FP)\cdot(TP+FN)\cdot(TN+FP)\cdot(TN+FN)}}$. The experimental growth and knockout data are extracted from (Kumelj et al., 2018). As a separate evaluation, we applied another method for identifying internal and unrealistic energy-generating cycles (Noor, 2018) , and no such cycles were found in Sco-GEM.

The simplest use of memote is generating snapshot reports showing the current state of the model. However, by integrating Travis CI [https://travis-ci.com/] into the gitHub repository, memote can be used to create a continuous report displaying how each commit affects the model quality. Memote version 0.9.12 was used in this work, and the memote snapshot report for Sco-GEM is given in the **Supplemental Information**.

#### Random sampling, normalization and pathway analysis

Because of the huge number of reactions in the EcSco-GEM, it is challenging to sample the solution space appropriately: we have chosen to use the method provided in the Raven Toolbox 2 (Bordel et al., 2010; Wang et al., 2018), which samples the vertices of the solution space. The drawback of this method is that it will not result in a uniform sampling of the solution space. However, it is more likely to span the entire solution space and also not prone to get stuck in extremely narrow parts of the solution space, which may happen with variants of the hit-and-run algorithm (Haraldsdóttir et al., 2017; Kaufman and Smith, 1998; Megchelenbrink et al., 2014). For each of the time points for each strain (17 different conditions in total) we constrained exchange reactions between 99% and 101% of the measured rates and generated 5000 random flux distributions with Gurobi as the solver. The reactions catalysed by isoenzymes were combined into the set of reactions in Sco-GEM and the reactions providing protein for each reaction. The mean of the 5000 flux distributions for each metabolic reaction was used in the following analysis.

Finally, for each of the 17 conditions, the mean fluxes were normalized by the $CO_2$ production rate. Then, the normalized mean fluxes were summarized for each metabolic pathway by using the curated pathway annotations, and we consider this a measure of the metabolic activity in each pathway. To ease visual interpretation of this data we used the function clustermap with default parameters in Seaborn version 0.9.0 (Michael Waskom et al., 2018) (which uses Scipy v1.3.1 (Virtanen et al., 2020)) to perform hierarchical clustering of pathways based on the metabolic activity in M145 (**Figure 2D**). We then kept this order in **Figure 3E** to enable strain comparison.

Since glucose and glutamate uptake rates, as well as growth rates were significantly different in the two strains and at different time points, normalization of the data was necessary to compare flux distributions. We tested various proxies as indicators of overall metabolic activity for normalization, namely $CO_2$ production; the total carbon uptake from glucose and glutamate; growth rate and mean flux value. As golden standard, we compared the fluxes through individual reactions that are well documented to change in M145 in response to the phosphate depletion (**Figure S9**). Normalization based on $CO_2$ production was tested and gave similar results than the data normalized on total carbon uptake from glucose and glutamate (**Figure S9A and S9B**. The data normalized by the sum of fluxes showed similar patterns as those achieved by glucose/glutamate and $CO_2$-normalized data but was noisier (**Figure S9C**). Considering the huge differences in growth rate, the growth-normalized data masked any other flux patterns (**Figure S9D**). The fact that different normalizations provided similar differences in metabolic fluxes proved that the inferred changes in metabolism were not artefacts of the normalization method but represent true metabolic activity of each strain.

#### Strains, cultivation conditions, sampling procedures, and analyses of media components and secondary metabolites

Experiments were performed using strain M145 of *S. coelicolor* A3(2) and its derivatives M1146 and M1152. The latter two are lacking the 4 major BGCs for actinorhodin (Act), undecylprodigiosin (Red), coelimycin P1 (Cpk), and calcium-dependent antibiotic (CDA), while M1152 is also carrying the pleiotropic, previously described antibiotic production enhancing mutation *rpoB* [S433L] (Gomez-Escribano and Bibb, 2011; Hu et al., 2002). All strains were kindly provided by Mervyn Bibb at John-Innes-Centre, Norwich, UK.

Triplicate cultivations of the strains were performed based on germinated spore inoculum on 1.8 L phosphate-limited medium SSBM-P, applying all routines of the optimized submerged batch fermentation strategy for *S. coelicolor* established and described before (Wentzel et al., 2012). All media were based on ion-free water, and all chemicals used were of analytical grade. In brief, spore batches of M145, M1146 and M1152 were generated by cultivation on soy flour-mannitol (SFM) agar plates (Kieser et al., 2000), harvesting by scraping off spores and suspension in 20% (v/v) glycerol, and storage in aliquots at −80 °C. $10^9$ CFU of spores of each strain were germinated for 5 hours at 30 °C and 250 rpm in 250 mL baffled shake-flasks with 2 g of 3 mm glass beads and 50 mL 2x YT medium (Claessen et al., 2003). The germinated spores were harvested by centrifugation (3200 x g, 15 °C, 5 min) and re-suspended in 5 mL ion-free water. An even dispersion of the germinated spores was achieved by vortex mixing (30 s), ensuring comparable inocula among biological replicas. Each bioreactor (1.8 liter starting volume culture medium in a 3-liter Applikon stirred tank reactor) was inoculated with 4.5 mL germinated spore suspension (corresponding to $9x10^8$ CFU). Phosphate-limited medium SSBM-P (Nieselt et al., 2010) consisted of Na-glutamate, 55.2 g/L; D-glucose, 40 g/L; $MgSO_4$, 2.0 mM; phosphate, 4.6 mM; supplemented minimal medium trace element solution SMM-TE (Claessen et al., 2003), 8 mL/L and TMS1, 5.6 mL/L. TMS1 consisted of $FeSO_4$ x 7 $H_2O$, 5 g/L; $CuSO_4$ x 5 $H_2O$, 390 mg/L; $ZnSO_4$ x 7 $H_2O$, 440 mg/L; $MnSO_4$ x $H_2O$, 150 mg/L; $Na_2MoO_4$ x 2 $H_2O$, 10 mg/L; $CoCl_2$ x 6 $H_2O$, 20 mg/L, and HCl, 50 mL/L. Clerol FBA 622 fermentation defoamer (Diamond Shamrock Scandinavia) was added to the growth medium before inoculation. Throughout fermentations, pH 7.0 was maintained constant by automatic addition of 2 M HCl. Dissolved oxygen levels were maintained at a minimum of 50% by automatic adjustment of the stirrer speed (minimal agitation 325 rpm).

The aeration rate was constant 0.5 L/(L x min) sterile air. Dissolved oxygen, agitation speed and carbon dioxide evolution rate were measured and logged on-line, while samples for the determination of cell dry weight, levels of growth medium components and secondary metabolites concentrations, as well as for transcriptome and proteome analysis were withdrawn throughout the fermentation trials as indicated in **Figure 2B**. For transcriptome analysis, 3 × 4 ml culture sample were applied in parallel onto three 0.45 µm nitrocellulose filters (Millipore) connected to vacuum. The biomass on each filter was immediately washed twice with 4 ml double-autoclaved ion-free water pre-heated to 30 °C, before the filters were collected in a 50 ml plastic tube, frozen in liquid nitrogen and stored at -80 °C until RNA isolation. For proteome analysis, 5 ml samples were taken and centrifuged (3200 x g, 5 min, 4 °C), and the resulting cell pellets frozen rapidly at -80 °C until further processing.

Levels of phosphate were measured spectrophotometrically by using the SpectroQuant Phosphate test kit (Merck KGaA, Darmstadt, Germany) following the manufacturer's instructions after downscaling to 96-well plate format. D-glucose and L-glutamate concentrations were determined by LC-MS using suitable standards, and measured concentrations were used to estimate specific uptake and excretion rates. Undecylprodigiosin (Red) levels were determined spectrophotometrically at 530 nm after acidified methanol extraction from the mycelium (Bystrykh et al., 1996). To determine relative amounts of actinorhodins (determined as total blue pigments, TBP), cell culture samples were treated with KOH (final concentration 1 M) and centrifuged, and the absorbance of the supernatants at 640 nm was determined (Bystrykh et al., 1996). Quantification of germicidin A and B was performed using targeted LC-MS analytics.

#### Proteomics

##### Sample preparation and NanoUPLC-MS analysis

Quantitative proteomics were performed using pipeline previously described (Gubbens et al., 2012). Mycelium pellets for proteome analysis were thawed and resuspended in the remaining liquid. 50 µL re-suspended mycelium was withdrawn and pelleted by centrifugation. 100 µL lysis buffer (4% SDS, 100 mM Tris-HCl pH 7.6, 50 mM EDTA) was added, and samples were sonicated in a water bath sonicator (Biorupter Plus, Diagenode) for 5 cycles of 30 s high power and 30 s off in ice water. Cell debris was pelleted and removed by centrifugation. Total protein was precipitated using the chloroform-methanol method described before (Wessel and Flügge, 1984). The pellet was dried in a vacuum centrifuge before dissolving in 0.1% RapiGest SF surfactant (Waters) at 95 °C. The protein concentration was measured at this stage using BCA method. Protein samples were then reduced by adding 5 mM DTT, followed by alkylation using 21.6 mM iodoacetamide. Then trypsin (recombinant, proteomics grade, Roche) was added at 0.1 µg per 10 µg protein. Samples were digested at 37 °C overnight. After digestion, trifluoroacetic acid was added to 0.5% followed by incubation at 37 °C for 30 min and centrifugation to remove MS interfering part of RapiGest SF. Peptide solution containing 8 µg peptide was then cleaned and desalted using STAGE-Tipping technique (Rappsilber et al., 2007). Final peptide concentration was adjusted to 40 ng/µL using sample solution (3% acetonitrile, 0.5% formic acid) for analysis.

200 ng (5 µL) digested peptide was injected and analysed by reversed-phase liquid chromatography on a nanoAcquity UPLC system (Waters) equipped with HSS-T3 C18 1.8 µm, 75 µm X 250 mm column (Waters). A gradient from 1% to 40% acetonitrile in 110 min (ending with a brief regeneration step to 90% for 3 min) was applied. [Glu$^{1}$]-fibrinopeptide B was used as lock mass compound and sampled every 30 s. Online MS/MS analysis was done using Synapt G2-Si HDMS mass spectrometer (Waters) with an UDMS$^{E}$ method set up as described in (Distler et al., 2014).

#### Data processing and label-free quantification

Raw data from all samples were first analysed using the vender software ProteinLynx Global SERVER (PLGS) version 3.0.3. Generally, mass spectrum data were generated using an MS$^{E}$ processing parameter with charge 2 lock mass 785.8426, and default energy thresholds. For protein identification, default workflow parameters except an additional acetyl in N-terminal variable modification were used. Reference protein database was downloaded from GenBank with the accession number NC_003888.3. The resulted dataset was imported to ISOQuant version 1.8 (Distler et al., 2014) for label-free quantification. Default high identification parameters were used in the quantification process. TOP3 result was converted to PPM (protein weight) and send to the modelers and others involved in interpreting the data (**Data Set S1, Tab 14**).

TOP3 quantification was filtered to remove identifications meet these two criteria: 1. identified in lower than 70% of samples of each strain and 2. sum of TOP3 value less than $1 \times 10^{5}$. Cleaned quantification data was further subjected to DESeq2 package version 1.22.2

(Love et al., 2014) and PCA was conducted after variance stabilizing transformation (vst) of normalized data.

#### Transcriptomics

##### RNA extraction and quality control

Bacteria were lysed using RNAprotect Bacteria (Qiagen) and following the manufacturer´s instruction. Briefly, filters containing bacteria were incubated with 4 ml of RNAprotect Bacteria reagent. After centrifugation, resulting samples were lysed using 500 µl of TE buffer (10 mM Tris·Cl, 1 mM EDTA, pH 8.0) containing 15 mg/ml lysozyme using 150-600 µm diameter glass beads (Sigma) agitated at 30 Hz for 5 minutes in the TissueLyser II (Qiagen).

Total RNA was extracted using RNeasy mini kit (Qiagen) and 700 µl of the resulting lysate complemented with 470 µl of absolute ethanol. RNAase-free DNase set (Qiagen) and centrifugation steps were performed to prevent DNA and ethanol contamination. Elution was performed using 30 µl of RNase-free water and by reloading the eluate on the column to improve the RNA yield. The RNA concentration was measured using Qubit RNA BR Assay Kit (ThermoFisher Scientific), RNA purity was assessed using A260/A280 and A260/A230 ratio using the Nano Drop ND-1000 Spectrophotometer (PEQLAB). RNA Integrity Number was estimated using RNA 6000 Nano Kit (Agilent) and the Bioanalyzer 2100 (Agilent).

##### Library preparation and sequencing

A total of 1 µg of total RNA was subjected to rRNA depletion using Ribo-Zero rRNA Removal Kit Bacteria (Illumina). The cDNA libraries were constructed using the resulting tRNA and the NEBNext Ultra II Directional RNA Library Prep Kit (NEB). Libraries were sequenced as single-reads (75 bp read length) on an Illumina NextSeq500 platform at a depth of 8–10 million reads each.

##### RNA-seq data assessment and analysis

Sequencing statistics including the quality per base and adapter content assessment of resulting transcriptome sequencing data were conducted with FastQC v0.11.5 (Andrews, 2016). All reads mappings were performed against the reference strain of *Streptomyces coelicolor A3(2)* (RefSeq ID NC_003888.3). The mappings of all samples were conducted with HISAT2 v2.1.0 (Kim et al., 2015). As parameters, spliced alignment of reads was disabled, and strand-specific information was set to reverse complemented (HISAT2 parameter --no-spliced-alignment and --rna-strandness "R"). The resulting mapping files in SAM format were converted to BAM format using SAMtools v1.6 (Li et al., 2009). Mapping statistics, including strand specificity estimation, percentage of mapped reads and fraction exonic region coverage, were conducted with the RNA-seq module of QualiMap2 v2.2.2-dev (Okonechnikov et al., 2016). Gene counts for all samples were computed with featureCounts v1.6.0 (Liao et al., 2014) based on the annotation of the respective reference genome, where the selected feature type was set to transcript records (featureCounts parameter -t transcript).

##### Normalization and differential gene expression

Raw count files were imported into Mayday SeaSight (Battke and Nieselt, 2011) for common, time-series-wide normalization. For this, the raw counts of all biological replicates of one strain across the time-series were log2-transformed (with pseudocount of +1 for the genes with zero counts) and then quantile-normalized. To make the two normalized time-series data of M154 and M1152 comparable, they were again quantile-normalized against each other. The normalized RNA-seq data are provided in **Data Set S1, Tab 15.**

Differentially expressed genes were identified by ANOVA using Orange (v3.2) and the bioinformatic toolkit (v), with FDR of $<0.01$ and a minimal fold enrichment $>1$ for at least one aligned time point. Genes with low expression ($\log_2 < 5$ for both strains and time points) were not considered for further analysis. The differentially expressed genes were subsequently scaled to the expression average and clustered by K-means. Visualization of genes and clusters were performed in python (v3.7) with matplotlib (v3.1.1). For this, the time-series of M145 and M1152 were aligned such that in the visual representation, the expression profiles of the two strains are aligned relative to the time point of phosphate depletion. Both DAVID (Huang et al., 2009a, 2009b) and the string database (Szklarczyk et al., 2019) was used to evaluate the function of each cluster, identifying overrepresentation of function groups based on GO annotation or text mining. Identified differential clusters or regulons were extracted from literature and plotted (**Data Set S1, Tab 5; Figure S8**). When we display the RNA-seq data as heatmaps (**Figure 6 and S3**) the order of genes is determined by hierarchical clustering using methods as previously described for the clustering of pathways based on metabolic activity.

### Estimation of growth, uptake and production rates for *Streptomyces coelicolor* M145 and M1152 from batch fermentation data

The estimated growth, uptake and secretion rates are based on average values of online and offline measurements of batch fermentation from three parallel bioreactors for each strain.

#### Growth rate estimation

Both the CDW (cell dry weight) measurements and $CO_2$ measurements can in principle be used to estimate growth rates, as there should be a linear relationship between the $CO_2$ concentration and cell mass. The $CO_2$ concentration is measured online on a high-resolution timescale (5 min) while CDW is measured offline with a four-hour resolution starting from 18 hour after inoculation.

To estimate growth rates, we have separated the growth into 5 different phases:

1. Lag phase - immediate phase after inoculation with no / low growth
2. Exponential growth - rapid growth after the initial lag phase
3. First linear growth rate - until phosphate depletion
4. Second linear growth rate – immediate phase after phosphate where there is still growth
5. Third linear growth rate – no or very low growth

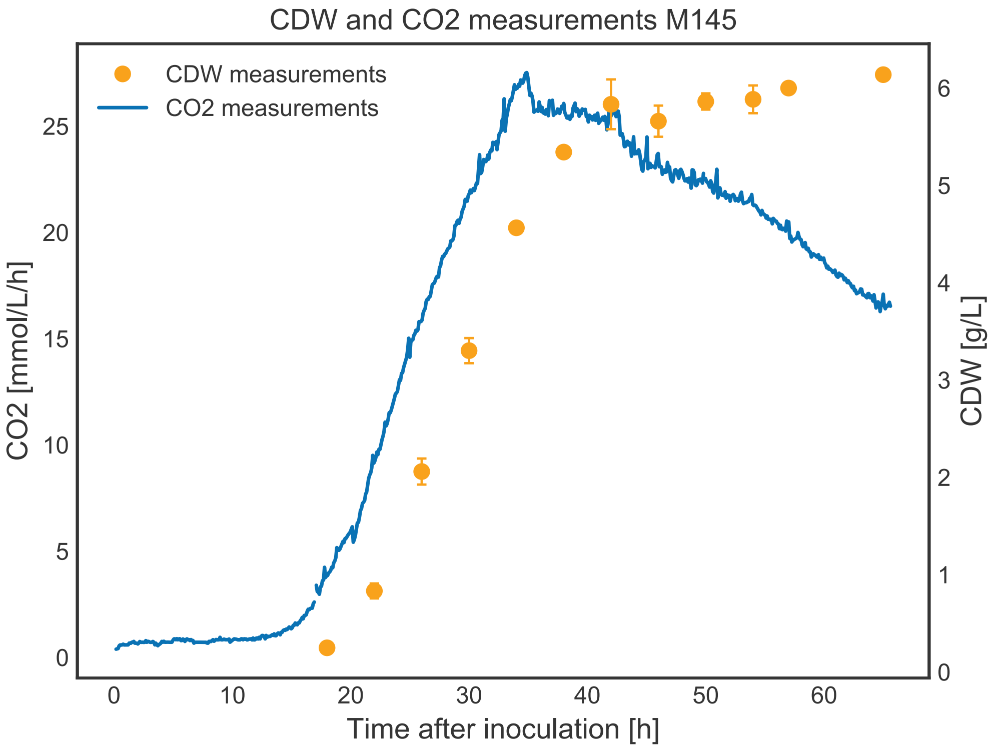


*Figure SX1: CDW and CO2 measurements of M145. We observe that the linear relationships between CDW and CO2 fails after 40 hours after inoculation.*

From Figure SX1 it is an obvious discrepancy between the CO2 curve and the CDW measurements after phosphate depletion (at 35 hours after inoculation for M145). Thus, despite the lower resolution we have decided to use the CDW measurements for the growth rate estimation except for the exponential growth phase.

#### Exponential growth rate from $CO_2$

The exponential growth rate was estimated by fitting an exponential curve on the form

$$X(t) = X_0 e^{\mu t}$$

to the selected region of the $CO_2$ measurements (Figure SX2 and Figure SX3) and given in Table SX1.

*Table SX1: Exponential growth rate estimated from $CO_2$-curves for M145 and M1152.*

| Strain | Estimated growth rate [$h^{-1}$] | Uncertainty [$h^{-1}$] |
|---|---|---|
| **M145** | 0.25 | 0.06 |
| **M1152** | 0.18 | 0.02 |

The uncertainty is estimated in a heuristic approach by observing the minimum and maximum values observed when changing the boundaries for the fitted function.

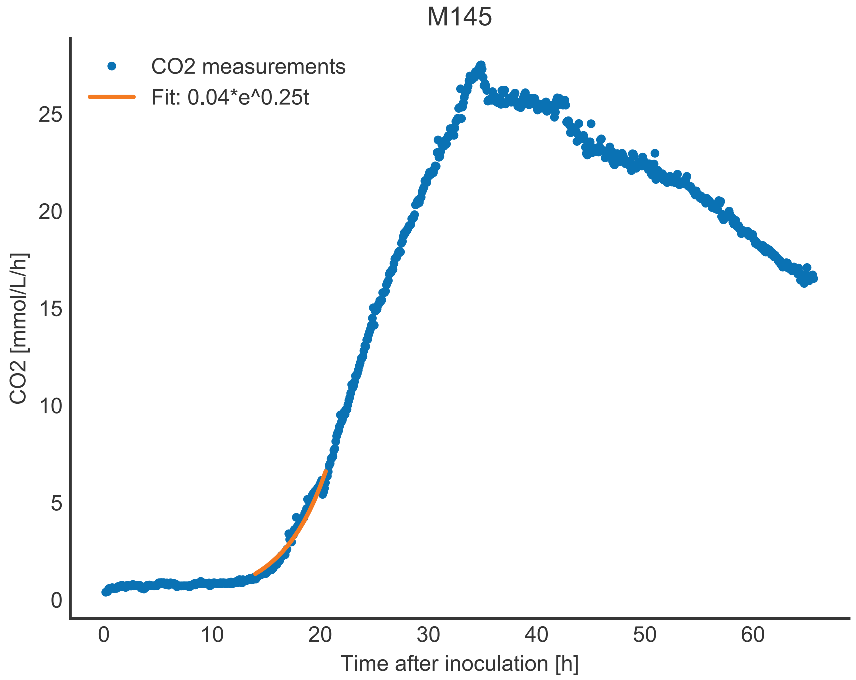


*Figure SX2: Fitting of exponential growth phase of CO2 measurements of M145.*

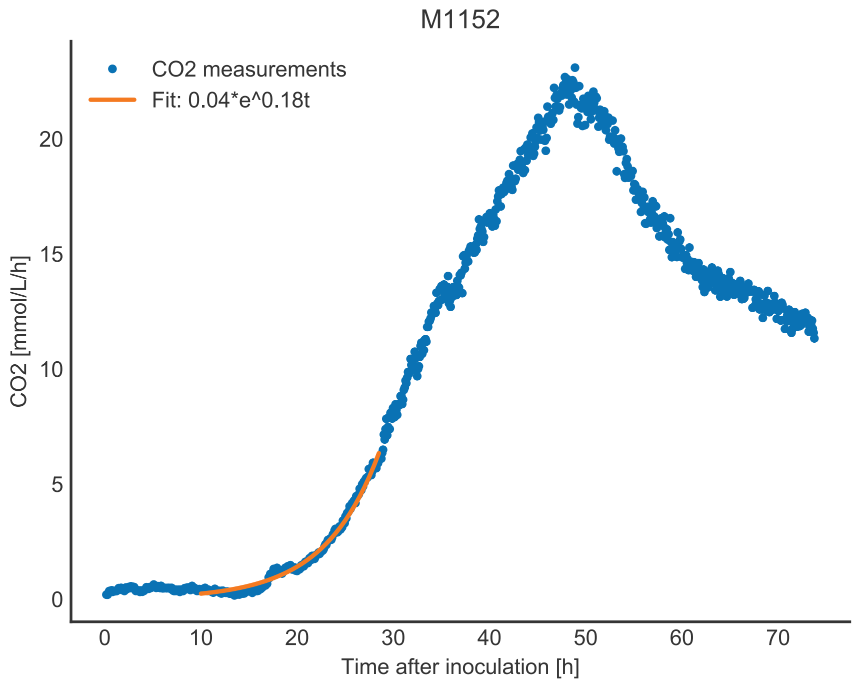


*Figure SX3: Fitting of exponential growth phase of CO2 measurements of M1152.*

#### Linear growth rates from CDW

The growth rate is estimated by fitting linear slopes to the three different linear phases of growth (Figure SX4 and Figure SX5). The specific growth rate is then calculated using the following equation

$$\frac{dX}{dt} = \mu X \;\rightarrow\; \mu = \frac{1}{X} \cdot \frac{dX}{dt}$$

where $\mu$ is the growth rate, X the CDW and $\frac{dX}{dt}$ is the slope of the linear fit. Because the inverse of the CDW the rates can become very large when the cell mass is low, but we use the estimated growth rate in the exponential phase as an upper bound. Predicted growth rates and CDW estimates at the timepoints for the proteome samples are given for M145 and M1152 in Table SX2 and Table SX3, respectively.

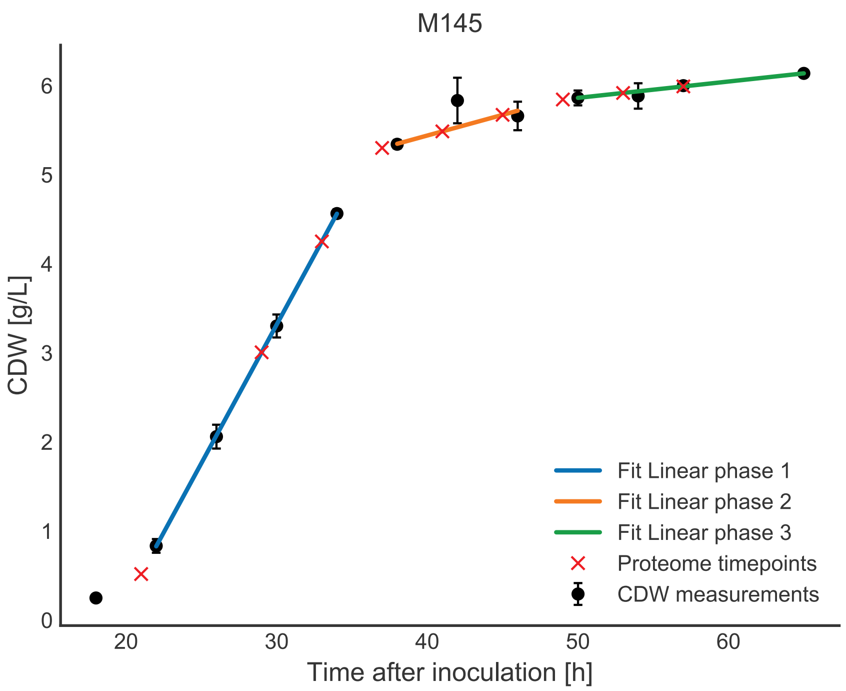


*Figure SX4: Piecewise linear fit of the CDW measurements of M145*

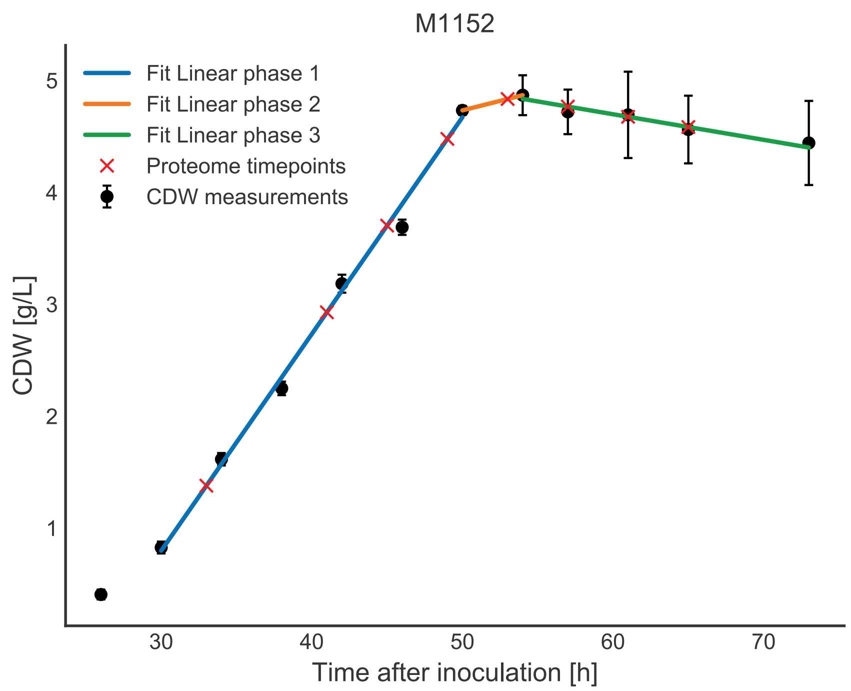


*Figure SX5: Piecewise linear fit of the CDW measurements of M1152*

#### Uptake rates of glucose and glutamic acid

The uptake rates for glucose and glutamate were also fitted using a piecewise linear function as seen in Figure SX6. Using the same time intervals as for the CDW estimates gave a very poor fit for M1152 and we therefore decided to use different time intervals. From the fitted slopes we estimated the uptake rates using the equation given below:

$$\frac{dS}{dt} = \mu_s X$$

where S is the substrate, $\mu_s$ the uptake rate and X the CDW at the given time. The uptake rates at 21 hours after inoculation seems to be too low for M145 and is caused by a too low estimate of the CDW. The uptake rates for glucose and given for M145 and M1152 in Table SX2 and Table SX3, respectively.

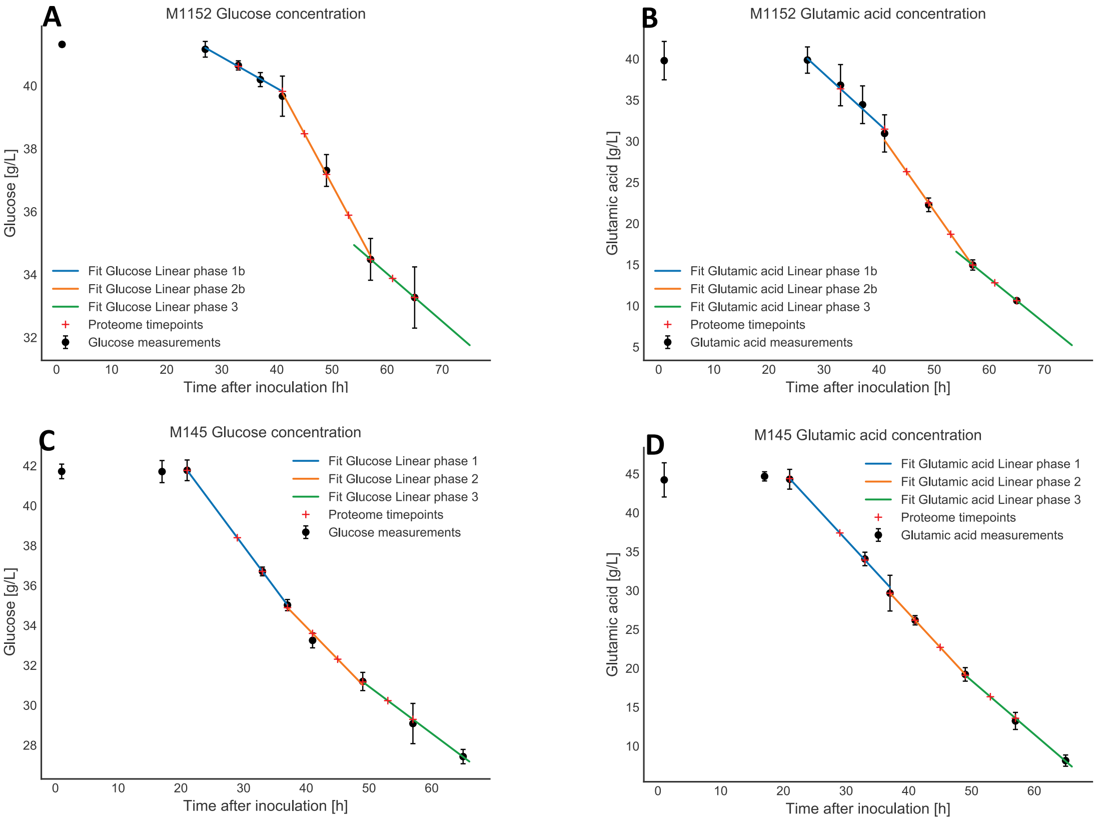


*Figure SX6: Fitted glucose and glutamic acid concentration for M1152 (A + B) and M145 (C + D). The intervals for M1152 were different than the intervals used for estimating the growth rate (A and B).*

#### Undecylprodigiosin (RED) production rate

We used the same method as for the uptake rates of glucose and glutamic acid to estimate the production rate of undecylprodigiosin (RED). The M1152 did not produce any RED (as expected). However, for the M145 the amount of undecylprodigiosin was measured both using MS (mass spectrometry) and OD (optical density). From the MS data it looks like the production of RED stops after approximately 53 hours, but from the OD measurements we observe a continuous increase until the end of the experiment. The hypothesis is that RED is continuously degraded into derivatives which are still measurable using OD but not using MS

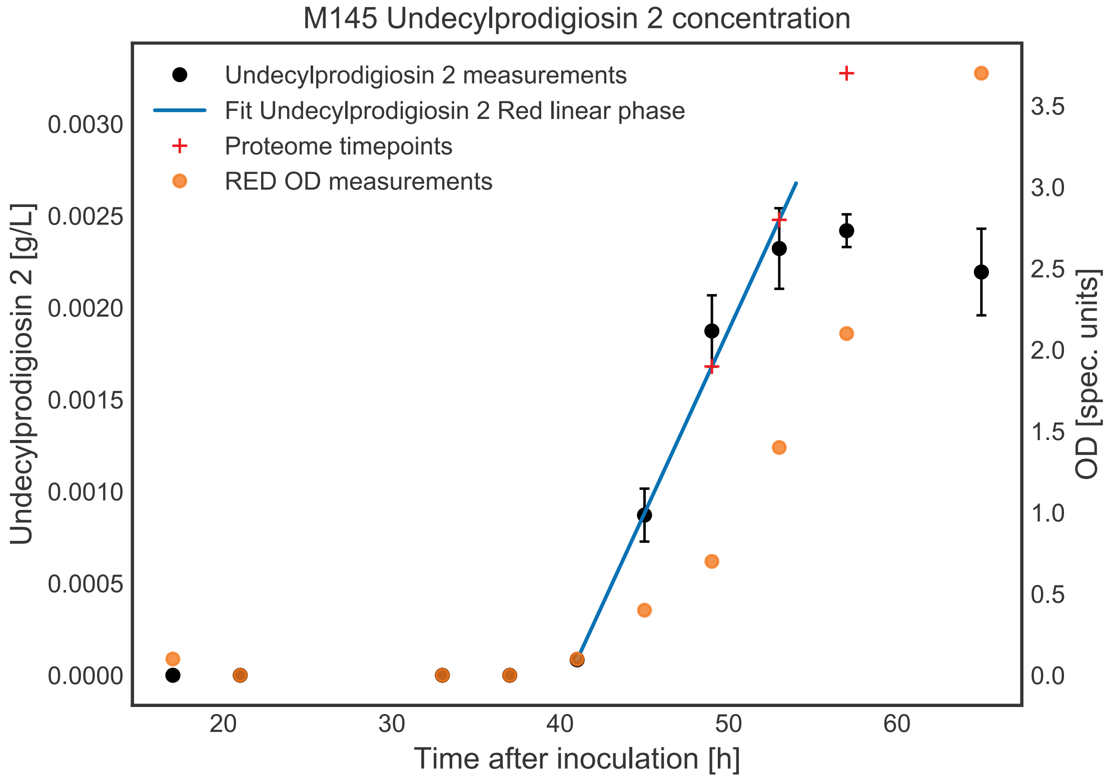


*Figure SX7: Estimated production rate of Undecylprodigiosin (RED).*

because of the different masses of the derivatives. Therefore, we have extrapolated the production rate of RED using the timepoints between 40 and 53 hours to estimate the production rate of RED at 57 hours (Figure SX7) .

#### Germicidin A and B production rates

We used the same method as for the uptake rates of glucose and glutamic acid to estimate the production rate of germicidin A and B for both M145 and M1152 (Figure SX8).

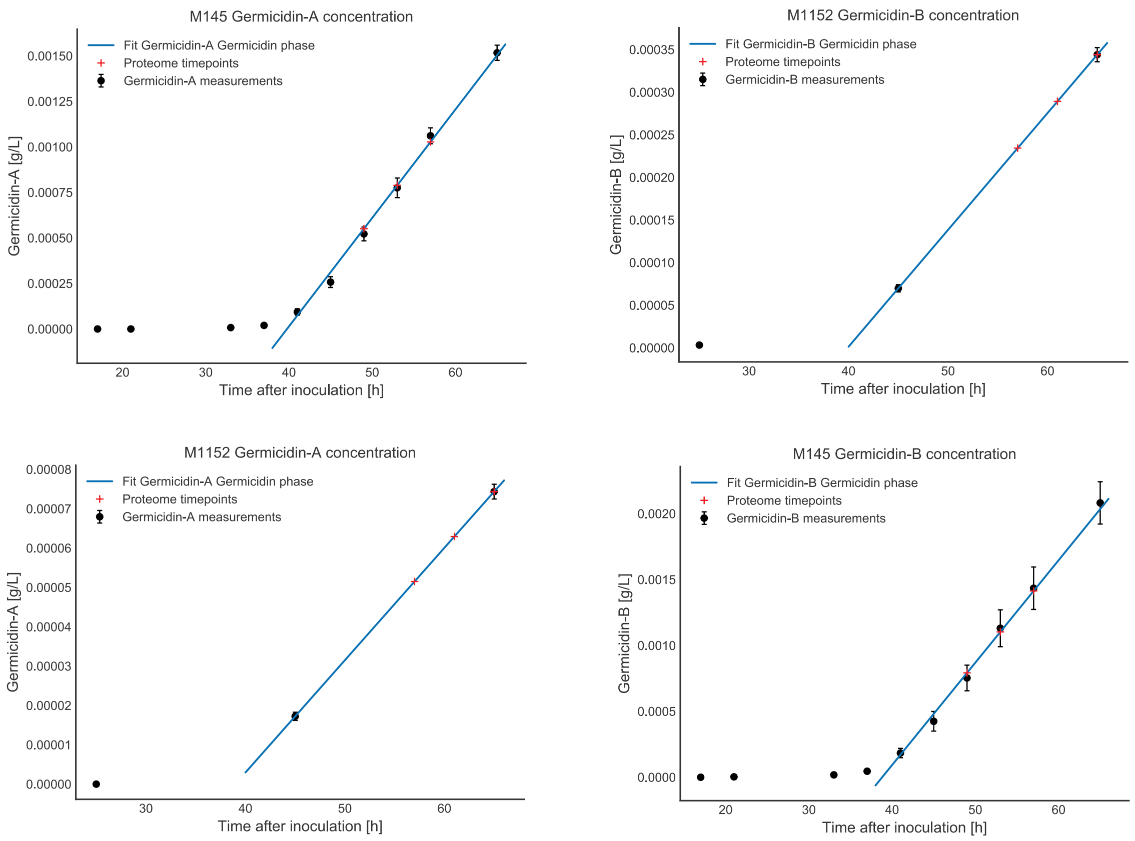


*Figure SX8: Estimation of production rates of germicidin A (left) and B (right) for M1152 and M145.*

*Table SX2: Estimated cell dry weight (CDW), growth rate and uptake / secretion rates for M145 at the timepoints for the proteome samples. The unit is mmol/g DW/h for the uptake / secretion rates.*

| TAI | Estimated CDW [g/L] | Growth rate $[h^{-1}]$ | Glucose | Glutamic acid | RED | Germicidin-A | Germicidin-B |
|---|---|---|---|---|---|---|---|
| **21** | 0.517 | 0.246* | -4.528[§] | -11.462[§] | 0 | 0 | 0 |
| **29** | 3.007 | 0.103 | -0.779 | -1.973 | 0 | 0 | 0 |
| **33** | 4.251 | 0.073 | -0.551 | -1.395 | 0 | 0 | 0 |
| **37** | 5.301 | 0.009 | -0.338 | -1.116 | 9.60E-05 | 5.70E-05 | 8.00E-05 |
| **41** | 5.487 | 0.008 | -0.327 | -1.078 | 9.20E-05 | 5.50E-05 | 7.80E-05 |
| **45** | 5.672 | 0.008 | -0.316 | -1.043 | 8.90E-05 | 5.40E-05 | 7.50E-05 |
| **49** | 5.845 | 0.003 | -0.223 | -0.803 | 8.70E-05 | 5.20E-05 | 7.30E-05 |

| | | | | | | | |
|---|---|---|---|---|---|---|---|
| **53** | 5.918 | 0.003 | -0.220 | -0.793 | 8.60E-05 | 5.10E-05 | 7.20E-05 |
| **57** | 5.991 | 0.003 | -0.217 | -0.784 | 8.50E-05 | 5.10E-05 | 7.10E-05 |

**This is the maximal growth rate predicted from the exponential fit of the $CO_2$ curve. The estimate rate from the linear fit of the CDW providedwas unrealisticly high.*

*§The values for the glucose and glutamate uptake rates are probably too high.*

*Table SX3: Estimated cell dry weight (CDW), growth rate and uptake / secretion rates for M1152 at the timepoints for the proteome samples. The unit is mmol/g DW/h for the uptake / secretion rates.*

| TAI | Estimated CDW [g/L] | Growth rate [$h^{-1}$] | Glucose | Glutamic acid | RED | Germicidin-A | Germicidin-B |
|---|---|---|---|---|---|---|---|
| **33** | 1.379 | 0.140 | -0.399 | -3.017 | 0 | 0 | 0 |
| **41** | 2.929 | 0.066 | -0.188 | -1.421 | 0 | 0 | 0 |
| **45** | 3.704 | 0.052 | -0.394 | -1.415 | 0 | 0 | 0 |
| **49** | 4.478 | 0.043 | -0.382 | -1.374 | 0 | 0 | 0 |
| **53** | 4.835 | 0.007 | -0.372 | -1.336 | 0 | 3.00E-06 | 1.56E-05 |
| **57** | 4.767 | -0.005 | -0.177 | -0.772 | 0 | 3.10E-06 | 1.58E-05 |
| **61** | 4.676 | -0.005 | -0.180 | -0.787 | 0 | 3.10E-06 | 1.61E-05 |
| **65** | 4.585 | -0.005 | -0.184 | -0.803 | 0 | 3.20E-06 | 1.64E-05 |

### Transparent methods references


Amara, A., Takano, E., Breitling, R., 2018. Development and validation of an updated computational model of Streptomyces coelicolor primary and secondary metabolism. BMC Genomics 19, 519. https://doi.org/10.1186/s12864-018-4905-5

Andrews, S., 2016. FastQC: a quality control tool for high throughput sequence data.

Bar-Even, A., Flamholz, A., Noor, E., Milo, R., 2012. Thermodynamic constraints shape the structure of carbon fixation pathways. Biochimica et Biophysica Acta (BBA) - Bioenergetics 1817, 1646–1659. https://doi.org/10.1016/j.bbabio.2012.05.002

Battke, F., Nieselt, K., 2011. Mayday SeaSight: combined analysis of deep sequencing and microarray data. PLoS ONE 6, e16345. https://doi.org/10.1371/journal.pone.0016345

Bordel, S., Agren, R., Nielsen, J., 2010. Sampling the Solution Space in Genome-Scale Metabolic Networks Reveals Transcriptional Regulation in Key Enzymes. PLOS Computational Biology 6, e1000859. https://doi.org/10.1371/journal.pcbi.1000859

Bystrykh, L.V., Fernández-Moreno, M.A., Herrema, J.K., Malpartida, F., Hopwood, D.A., Dijkhuizen, L., 1996. Production of actinorhodin-related “blue pigments” by Streptomyces coelicolor A3(2). J. Bacteriol. 178, 2238–2244.

Caspi, R., Altman, T., Billington, R., Dreher, K., Foerster, H., Fulcher, C.A., Holland, T.A., Keseler, I.M., Kothari, A., Kubo, A., Krummenacker, M., Latendresse, M., Mueller, L.A., Ong, Q., Paley, S., Subhraveti, P., Weaver, D.S., Weerasinghe, D., Zhang, P., Karp, P.D., 2014. The MetaCyc database of metabolic pathways and enzymes and the BioCyc collection of Pathway/Genome Databases. Nucleic Acids Res 42, D459–D471. https://doi.org/10.1093/nar/gkt1103

Chicco, D., Jurman, G., 2020. The advantages of the Matthews correlation coefficient (MCC) over F1 score and accuracy in binary classification evaluation. BMC Genomics 21, 6. https://doi.org/10.1186/s12864-019-6413-7

Claessen, D., Rink, R., Jong, W. de, Siebring, J., Vreugd, P. de, Boersma, F.G.H., Dijkhuizen, L., Wösten, H.A.B., 2003. A novel class of secreted hydrophobic proteins is involved in aerial hyphae formation in Streptomyces coelicolor by forming amyloid-like fibrils. Genes Dev. 17, 1714–1726. https://doi.org/10.1101/gad.264303

Cokelaer, T., Pultz, D., Harder, L.M., Serra-Musach, J., Saez-Rodriguez, J., 2013. BioServices: a common Python package to access biological Web Services programmatically. Bioinformatics 29, 3241–3242. https://doi.org/10.1093/bioinformatics/btt547

Courtot, M., Juty, N., Knüpfer, C., Waltemath, D., Zhukova, A., Dräger, A., Dumontier, M., Finney, A., Golebiewski, M., Hastings, J., Hoops, S., Keating, S., Kell, D.B., Kerrien, S., Lawson, J., Lister, A., Lu, J., Machne, R., Mendes, P., Pocock, M., Rodriguez, N., Villeger, A., Wilkinson, D.J., Wimalaratne, S., Laibe, C., Hucka, M., Le Novère, N., 2011. Controlled vocabularies and semantics in systems biology. Mol. Syst. Biol. 7, 543. https://doi.org/10.1038/msb.2011.77

Distler, U., Kuharev, J., Navarro, P., Levin, Y., Schild, H., Tenzer, S., 2014. Drift time-specific collision energies enable deep-coverage data-independent acquisition proteomics. Nat. Methods 11, 167–170. https://doi.org/10.1038/nmeth.2767

Ebrahim, A., Lerman, J.A., Palsson, B.O., Hyduke, D.R., 2013. COBRApy: COnstraints-Based Reconstruction and Analysis for Python. BMC Systems Biology 7, 74. https://doi.org/10.1186/1752-0509-7-74

Elbourne, L.D.H., Tetu, S.G., Hassan, K.A., Paulsen, I.T., 2017. TransportDB 2.0: a database for exploring membrane transporters in sequenced genomes from all domains of life. Nucleic Acids Research 45, D320–D324. https://doi.org/10.1093/nar/gkw1068

Feist, A.M., Henry, C.S., Reed, J.L., Krummenacker, M., Joyce, A.R., Karp, P.D., Broadbelt, L.J., Hatzimanikatis, V., Palsson, B.Ø., 2007. A genome-scale metabolic reconstruction for Escherichia coli K-12 MG1655 that accounts for 1260 ORFs and thermodynamic information. Molecular Systems Biology 3, 121. https://doi.org/10.1038/msb4100155

Flamholz, A., Noor, E., Bar-Even, A., Milo, R., 2012. eQuilibrator—the biochemical thermodynamics calculator. Nucleic Acids Res 40, D770–D775. https://doi.org/10.1093/nar/gkr874

Fritzemeier, C.J., Hartleb, D., Szappanos, B., Papp, B., Lercher, M.J., 2017. Erroneous energy-generating cycles in published genome scale metabolic networks: Identification and removal. PLOS Computational Biology 13, e1005494. https://doi.org/10.1371/journal.pcbi.1005494

Gomez-Escribano, J.P., Bibb, M.J., 2011. Engineering Streptomyces coelicolor for heterologous expression of secondary metabolite gene clusters. Microbial Biotechnology 4, 207–215. https://doi.org/10.1111/j.1751-7915.2010.00219.x

Gubbens, J., Janus, M., Florea, B.I., Overkleeft, H.S., van Wezel, G.P., 2012. Identification of glucose kinase-dependent and -independent pathways for carbon control of primary metabolism, development and antibiotic production in *Streptomyces coelicolor* by quantitative proteomics. Molecular Microbiology 86, 1490–1507. https://doi.org/10.1111/mmi.12072

Haraldsdóttir, H.S., Cousins, B., Thiele, I., Fleming, R.M.T., Vempala, S., 2017. CHRR: coordinate hit-and-run with rounding for uniform sampling of constraint-based models. Bioinformatics 33, 1741–1743. https://doi.org/10.1093/bioinformatics/btx052

Hu, H., Zhang, Q., Ochi, K., 2002. Activation of Antibiotic Biosynthesis by Specified Mutations in the rpoB Gene (Encoding the RNA Polymerase β Subunit) of Streptomyces lividans.

Journal of Bacteriology 184, 3984–3991. https://doi.org/10.1128/JB.184.14.3984-3991.2002

Huang, D.W., Sherman, B.T., Lempicki, R.A., 2009a. Bioinformatics enrichment tools: paths toward the comprehensive functional analysis of large gene lists. Nucleic Acids Res. 37, 1–13. https://doi.org/10.1093/nar/gkn923

Huang, D.W., Sherman, B.T., Lempicki, R.A., 2009b. Systematic and integrative analysis of large gene lists using DAVID bioinformatics resources. Nat Protoc 4, 44–57. https://doi.org/10.1038/nprot.2008.211

Jeske, L., Placzek, S., Schomburg, I., Chang, A., Schomburg, D., 2019. BRENDA in 2019: a European ELIXIR core data resource. Nucleic Acids Res 47, D542–D549. https://doi.org/10.1093/nar/gky1048

Kanehisa, M., 2000. KEGG: Kyoto Encyclopedia of Genes and Genomes. Nucleic Acids Research 28, 27–30. https://doi.org/10.1093/nar/28.1.27

Kanehisa, M., Sato, Y., Furumichi, M., Morishima, K., Tanabe, M., 2019. New approach for understanding genome variations in KEGG. Nucleic Acids Research 47, D590–D595. https://doi.org/10.1093/nar/gky962

Karp, P.D., Billington, R., Caspi, R., Fulcher, C.A., Latendresse, M., Kothari, A., Keseler, I.M., Krummenacker, M., Midford, P.E., Ong, Q., Ong, W.K., Paley, S.M., Subhraveti, P., 2017. The BioCyc collection of microbial genomes and metabolic pathways. Brief Bioinform. https://doi.org/10.1093/bib/bbx085

Karp, P.D., Latendresse, M., Paley, S.M., Krummenacker, M., Ong, Q.D., Billington, R., Kothari, A., Weaver, D., Lee, T., Subhraveti, P., Spaulding, A., Fulcher, C., Keseler, I.M., Caspi, R., 2016. Pathway Tools version 19.0 update: software for pathway/genome informatics and systems biology. Brief Bioinform 17, 877–890. https://doi.org/10.1093/bib/bbv079

Kaufman, D.E., Smith, R.L., 1998. Direction Choice for Accelerated Convergence in Hit-and-Run Sampling. Operations Research 46, 84–95. https://doi.org/10.1287/opre.46.1.84

Keesey, J.K., Bigelis, R., Fink, G.R., 1979. The product of the his4 gene cluster in Saccharomyces cerevisiae. A trifunctional polypeptide. J. Biol. Chem. 254, 7427–7433.

Kieser, T., Bibb, M.J., Buttner, M.J., Chater, K.F., Hopwood, D.A., 2000. Practical Streptomyces Genetics. John Innes Foundation, Norwich, UK.

Kim, D., Langmead, B., Salzberg, S.L., 2015. HISAT: a fast spliced aligner with low memory requirements. Nat. Methods 12, 357–360. https://doi.org/10.1038/nmeth.3317

King, Z.A., Lu, J., Dräger, A., Miller, P., Federowicz, S., Lerman, J.A., Ebrahim, A., Palsson, B.O., Lewis, N.E., J., H., 2016. BiGG Models: A platform for integrating, standardizing and sharing genome-scale models. Nucleic Acids Research 44, D515–D522. https://doi.org/10.1093/nar/gkv1049

Kumelj, T., Sulheim, S., Wentzel, A., Almaas, E., 2018. Predicting Strain Engineering Strategies Using iKS1317: A Genome-Scale Metabolic Model of Streptomyces coelicolor. Biotechnology Journal 0, 1800180. https://doi.org/10.1002/biot.201800180

Li, H., Handsaker, B., Wysoker, A., Fennell, T., Ruan, J., Homer, N., Marth, G., Abecasis, G., Durbin, R., 1000 Genome Project Data Processing Subgroup, 2009. The Sequence Alignment/Map format and SAMtools. Bioinformatics 25, 2078–2079. https://doi.org/10.1093/bioinformatics/btp352

Liao, Y., Smyth, G.K., Shi, W., 2014. featureCounts: an efficient general purpose program for assigning sequence reads to genomic features. Bioinformatics 30, 923–930. https://doi.org/10.1093/bioinformatics/btt656

Lieven, C., Beber, M.E., Olivier, B.G., Bergmann, F.T., Ataman, M., Babaei, P., Bartell, J.A., Blank, L.M., Chauhan, S., Correia, K., Diener, C., Dräger, A., Ebert, B.E., Edirisinghe, J.N., Faria, J.P., Feist, A., Fengos, G., Fleming, R.M.T., Garcia-Jimenez, B., Hatzimanikatis, V., Helvoirt, W. van, Henry, C., Hermjakob, H., Herrgard, M.J., Kim, H.U., King, Z., Koehorst, J.J., Klamt, S., Klipp, E., Lakshmanan, M., Novere, N.L., Lee, D.-Y., Lee, S.Y., Lee, S., Lewis, N.E., Ma, H., Machado, D., Mahadevan, R., Maia, P., Mardinoglu, A., Medlock, G.L., Monk, J., Nielsen, J., Nielsen, L.K., Nogales, J., Nookaew, I., Resendis, O., Palsson, B., Papin, J.A., Patil, K.R., Poolman, M., Price, N.D., Richelle, A., Rocha, I., Sanchez, B., Schaap, P., Sheriff, R.S.M., Shoaie, S., Sonnenschein, N., Teusink, B., Vilaca, P., Vik, J.O., Wodke, J.A., Xavier, J.C., Yuan, Q., Zakhartsev, M., Zhang, C., 2018. Memote: A community-driven effort towards a standardized genome-scale metabolic model test suite. bioRxiv 350991. https://doi.org/10.1101/350991

Love, M.I., Huber, W., Anders, S., 2014. Moderated estimation of fold change and  dispersion for RNA-seq data with DESeq2. Genome Biol. 15, 550. https://doi.org/10.1186/s13059-014-0550-8

Maurer, K.H., Pfeiffer, F., Zehender, H., Mecke, D., 1983. Characterization of two glyceraldehyde-3-phosphate dehydrogenase isoenzymes from the pentalenolactone producer Streptomyces arenae. J. Bacteriol. 153, 930–936.

Megchelenbrink, W., Huynen, M., Marchiori, E., 2014. optGpSampler: An Improved Tool for Uniformly Sampling the Solution-Space of Genome-Scale Metabolic Networks. PLOS ONE 9, e86587. https://doi.org/10.1371/journal.pone.0086587

Michael Waskom, Olga Botvinnik, Drew O'Kane, Paul Hobson, Joel Ostblom, Saulius Lukauskas, David C Gemperline, Tom Augspurger, Yaroslav Halchenko, John B. Cole, Jordi Warmenhoven, Julian de Ruiter, Cameron Pye, Stephan Hoyer, Jake Vanderplas, Santi Villalba, Gero Kunter, Eric Quintero, Pete Bachant, Marcel Martin, Kyle Meyer, Alistair Miles, Yoav Ram, Thomas Brunner, Tal Yarkoni, Mike Lee Williams, Constantine Evans, Clark Fitzgerald, Brian, Adel Qalieh, 2018. mwaskom/seaborn: v0.9.0 (July 2018). Zenodo. https://doi.org/10.5281/zenodo.1313201

Moretti, S., Martin, O., Van Du Tran, T., Bridge, A., Morgat, A., Pagni, M., 2016. MetaNetX/MNXref – reconciliation of metabolites and biochemical reactions to bring together genome-scale metabolic networks. Nucleic Acids Res 44, D523–D526. https://doi.org/10.1093/nar/gkv1117

NCBI Resource Coordinators, 2017. Database Resources of the National Center for Biotechnology Information. Nucleic Acids Research 45, D12–D17. https://doi.org/10.1093/nar/gkw1071

Nieselt, K., Battke, F., Herbig, A., Bruheim, P., Wentzel, A., Jakobsen, Ø.M., Sletta, H., Alam, M.T., Merlo, M.E., Moore, J., Omara, W.A., Morrissey, E.R., Juarez-Hermosillo, M.A., Rodríguez-García, A., Nentwich, M., Thomas, L., Iqbal, M., Legaie, R., Gaze, W.H., Challis, G.L., Jansen, R.C., Dijkhuizen, L., Rand, D.A., Wild, D.L., Bonin, M., Reuther, J., Wohlleben, W., Smith, M.C., Burroughs, N.J., Martín, J.F., Hodgson, D.A., Takano, E., Breitling, R., Ellingsen, T.E., Wellington, E.M., 2010. The dynamic architecture of the metabolic switch in Streptomyces coelicolor. BMC Genomics 11, 10. https://doi.org/10.1186/1471-2164-11-10

Noor, E., 2018. Removing both Internal and Unrealistic Energy-Generating Cycles in Flux Balance Analysis. arXiv:1803.04999 [q-bio].

Noor, E., Haraldsdóttir, H.S., Milo, R., Fleming, R.M.T., 2013. Consistent Estimation of Gibbs Energy Using Component Contributions. PLOS Computational Biology 9, e1003098. https://doi.org/10.1371/journal.pcbi.1003098

Okonechnikov, K., Conesa, A., García-Alcalde, F., 2016. Qualimap 2: advanced multi-sample quality control for high-throughput sequencing data. Bioinformatics 32, 292–294. https://doi.org/10.1093/bioinformatics/btv566

Rappsilber, J., Mann, M., Ishihama, Y., 2007. Protocol for micro-purification, enrichment, pre-fractionation and storage of peptides for proteomics using StageTips. Nat Protoc 2, 1896–1906. https://doi.org/10.1038/nprot.2007.261

Rauch, G., Ehammer, H., Bornemann, S., Macheroux, P., 2008. Replacement of two invariant serine residues in chorismate synthase provides evidence that a proton relay system is essential for intermediate formation and catalytic activity: Proton relay system in chorismate synthase. FEBS Journal 275, 1464–1473. https://doi.org/10.1111/j.1742-4658.2008.06305.x

Saier, M.H., Reddy, V.S., Tsu, B.V., Ahmed, M.S., Li, C., Moreno-Hagelsieb, G., 2016. The Transporter Classification Database (TCDB): recent advances. Nucleic Acids Res 44, D372–D379. https://doi.org/10.1093/nar/gkv1103

Sánchez, B.J., Zhang, C., Nilsson, A., Lahtvee, P.-J., Kerkhoven, E.J., Nielsen, J., 2017. Improving the phenotype predictions of a yeast genome-scale metabolic model by incorporating enzymatic constraints. Molecular Systems Biology 13, 935. https://doi.org/10.15252/msb.20167411

Schendel, F.J., Mueller, E., Stubbe, J., Shiau, A., Smith, J.M., 1989. Formylglycinamide ribonucleotide synthetase from Escherichia coli: cloning, sequencing, overproduction, isolation, and characterization. Biochemistry 28, 2459–2471. https://doi.org/10.1021/bi00432a017

Szklarczyk, D., Gable, A.L., Lyon, D., Junge, A., Wyder, S., Huerta-Cepas, J., Simonovic, M., Doncheva, N.T., Morris, J.H., Bork, P., Jensen, L.J., Mering, C. von, 2019. STRING v11: protein-protein association networks with increased coverage, supporting functional discovery in genome-wide experimental datasets. Nucleic Acids Res. 47, D607–D613. https://doi.org/10.1093/nar/gky1131

Talfournier, F., Stines-Chaumeil, C., Branlant, G., 2011. Methylmalonate semialdehyde dehydrogenase from Bacillus subtilis : substrate specificity and coenzyme A binding. J. Biol. Chem. jbc.M110.213280. https://doi.org/10.1074/jbc.M110.213280

The UniProt Consortium, 2019. UniProt: a worldwide hub of protein knowledge. Nucleic Acids Res 47, D506–D515. https://doi.org/10.1093/nar/gky1049

Thiele, I., Palsson, B.Ø., 2010. A protocol for generating a high-quality genome-scale metabolic reconstruction. Nature protocols 5, 93–121. https://doi.org/10.1038/nprot.2009.203

Virtanen, P., Gommers, R., Oliphant, T.E., Haberland, M., Reddy, T., Cournapeau, D., Burovski, E., Peterson, P., Weckesser, W., Bright, J., Walt, S.J. van der, Brett, M., Wilson, J., Millman, K.J., Mayorov, N., Nelson, A.R.J., Jones, E., Kern, R., Larson, E., Carey, C.J., Polat, İ., Feng, Y., Moore, E.W., VanderPlas, J., Laxalde, D., Perktold, J., Cimrman, R., Henriksen, I., Quintero, E.A., Harris, C.R., Archibald, A.M., Ribeiro, A.H., Pedregosa, F., Mulbregt, P. van, 2020. SciPy 1.0: fundamental algorithms for scientific computing in Python. Nat Methods 17, 261–272. https://doi.org/10.1038/s41592-019-0686-2

Wang, H., Marcišauskas, S., Sánchez, B.J., Domenzain, I., Hermansson, D., Agren, R., Nielsen, J., Kerkhoven, E.J., 2018. RAVEN 2.0: A versatile toolbox for metabolic network reconstruction and a case study on Streptomyces coelicolor. PLOS Computational Biology 14, e1006541. https://doi.org/10.1371/journal.pcbi.1006541

Wentzel, A., Bruheim, P., Øverby, A., Jakobsen, Ø.M., Sletta, H., Omara, W.A.M., Hodgson, D.A., Ellingsen, T.E., 2012. Optimized submerged batch fermentation strategy for systems scale studies of metabolic switching in Streptomyces coelicolor A3(2). BMC systems biology 6, 59. https://doi.org/10.1186/1752-0509-6-59

Wessel, D., Flügge, U.I., 1984. A method for the quantitative recovery of protein in dilute solution in the presence of detergents and lipids. Anal. Biochem. 138, 141–143.
