## Supplementary figures and images for "Enzyme-constrained models and omics analysis of Streptomyces coelicolor reveal metabolic changes that enhance heterologous production"

### Supplemental figures

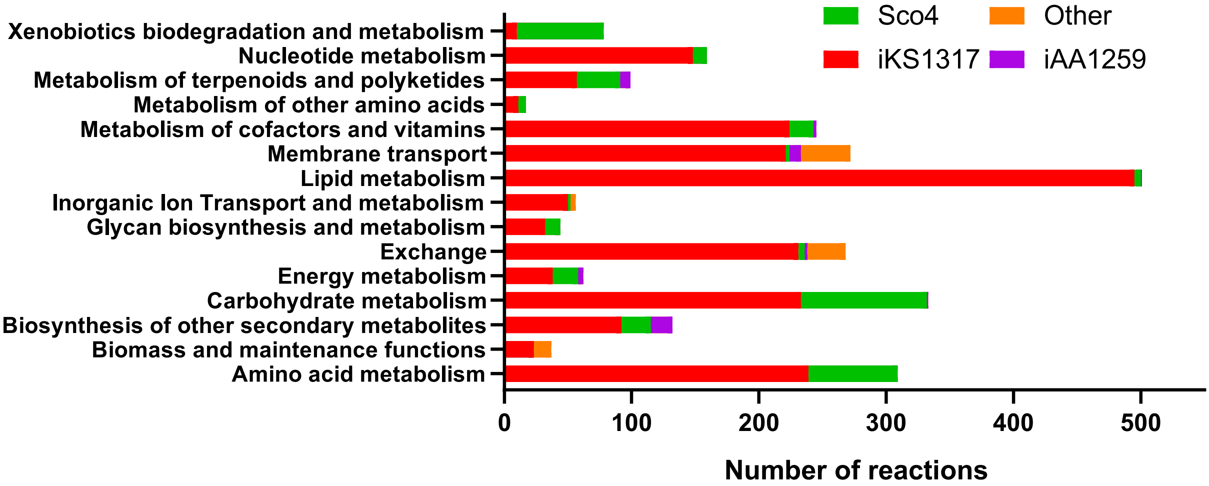

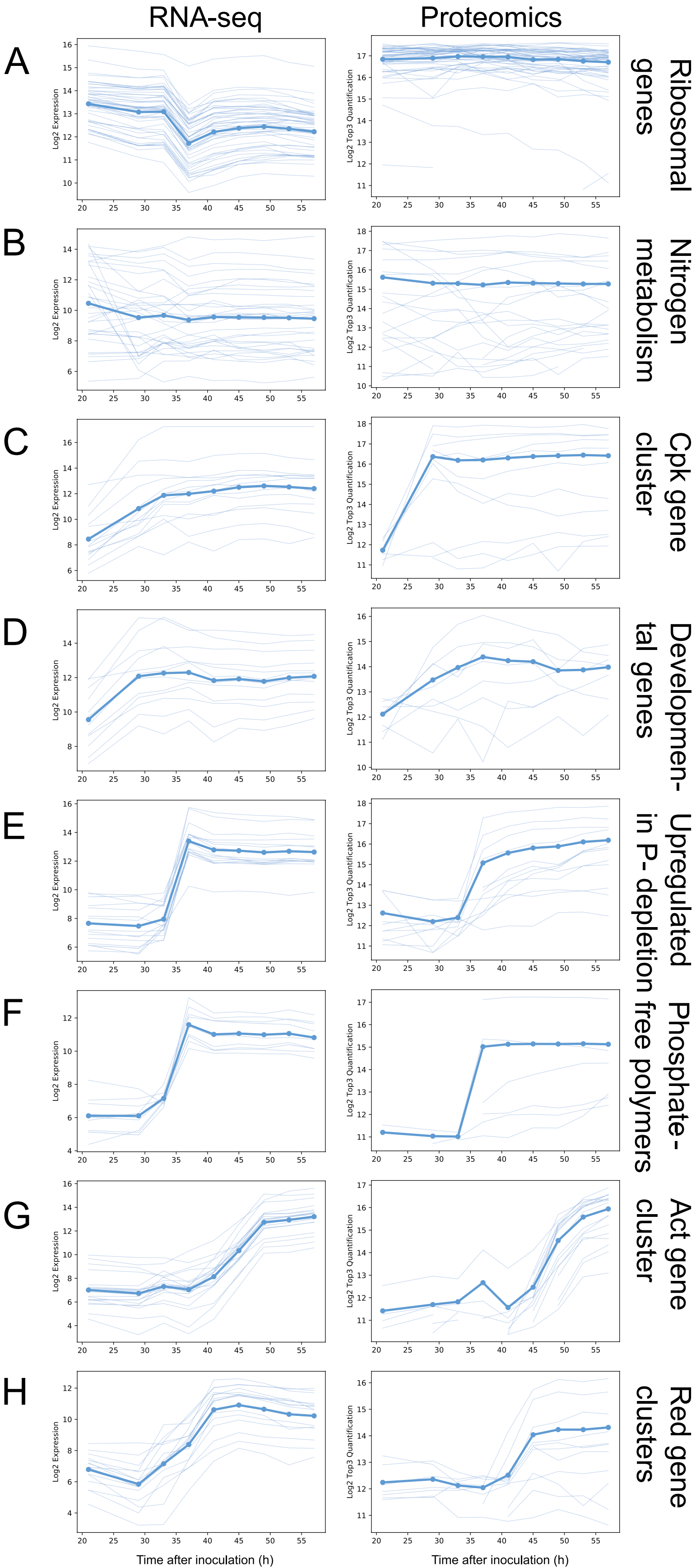

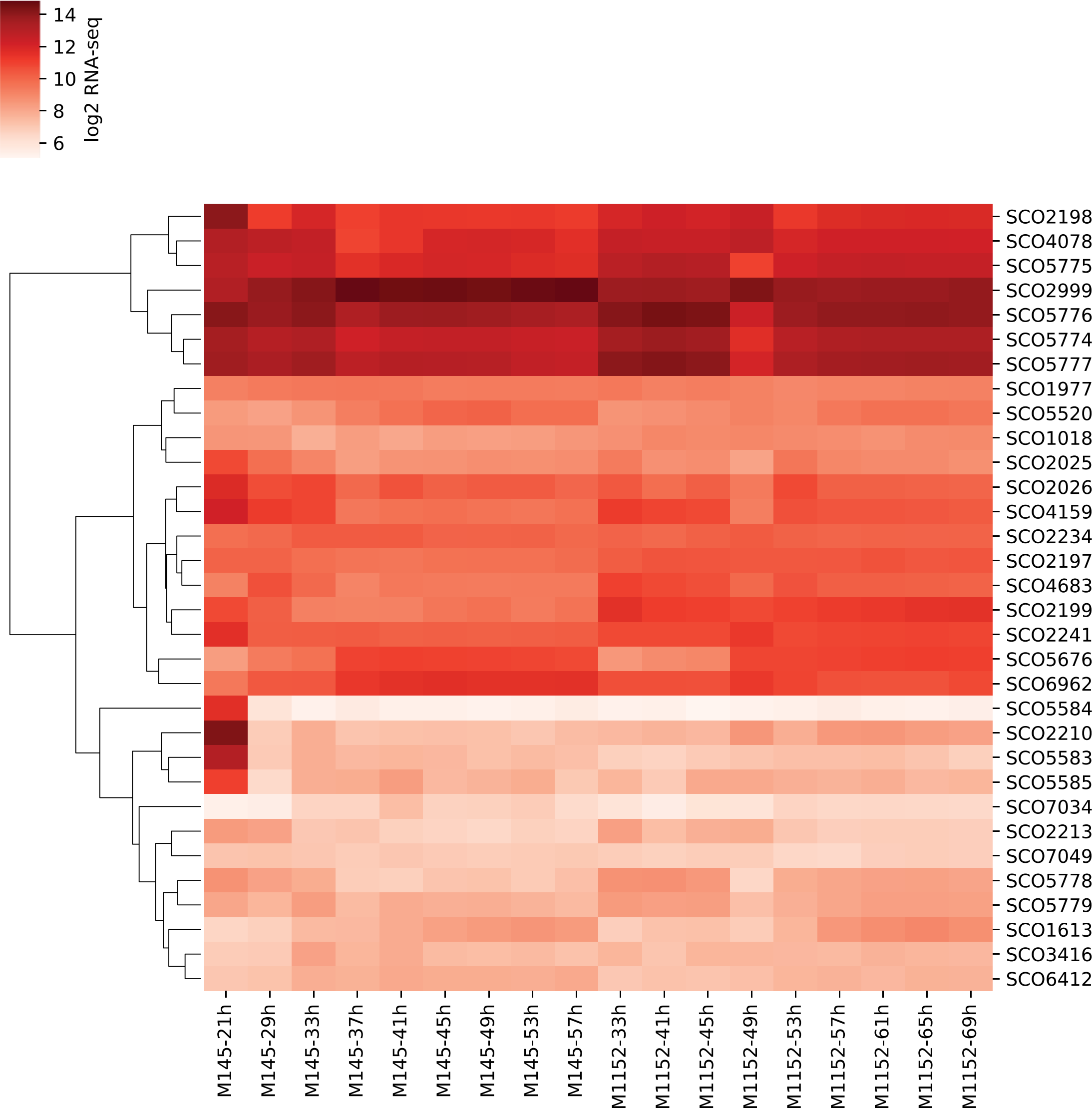

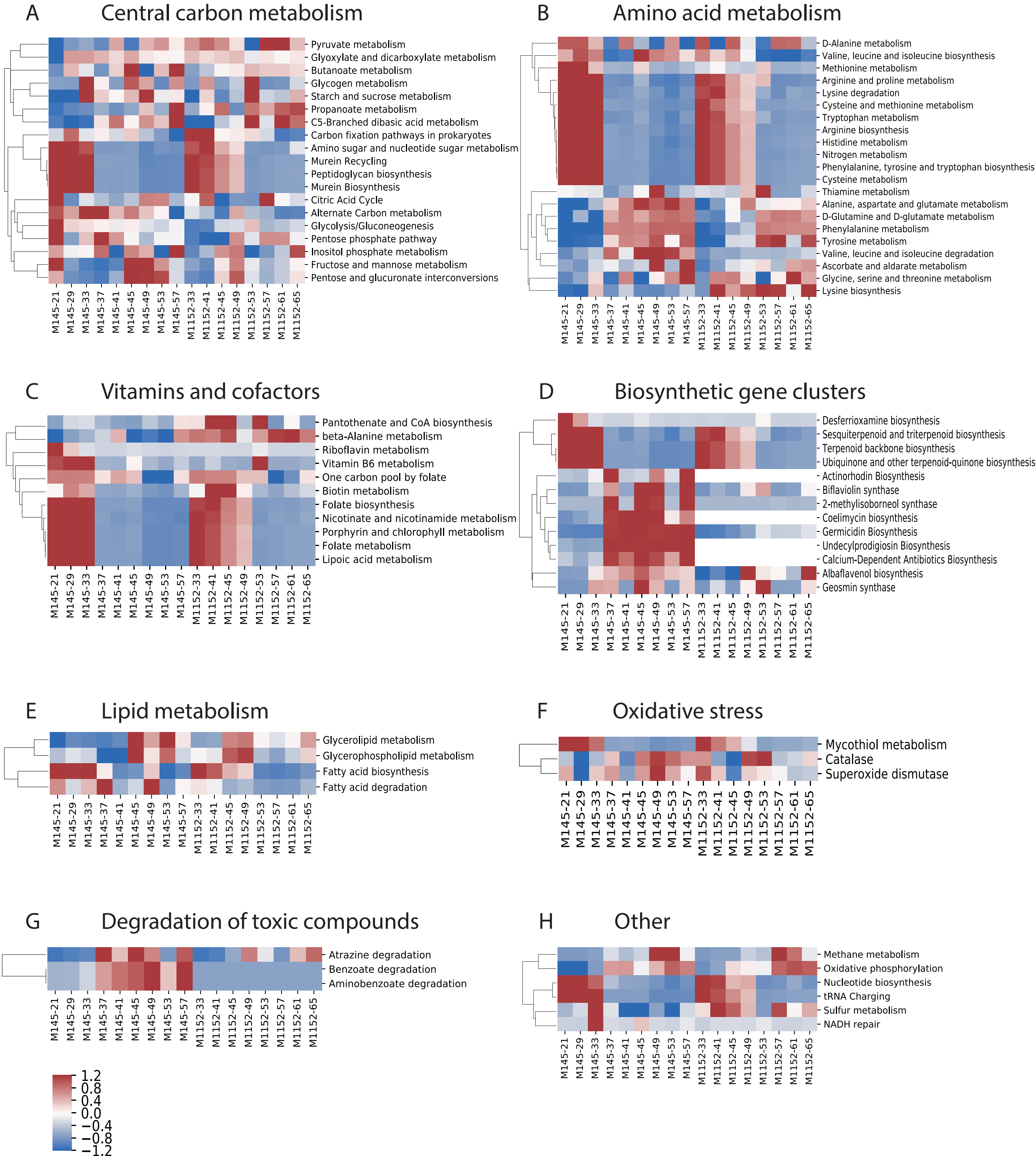

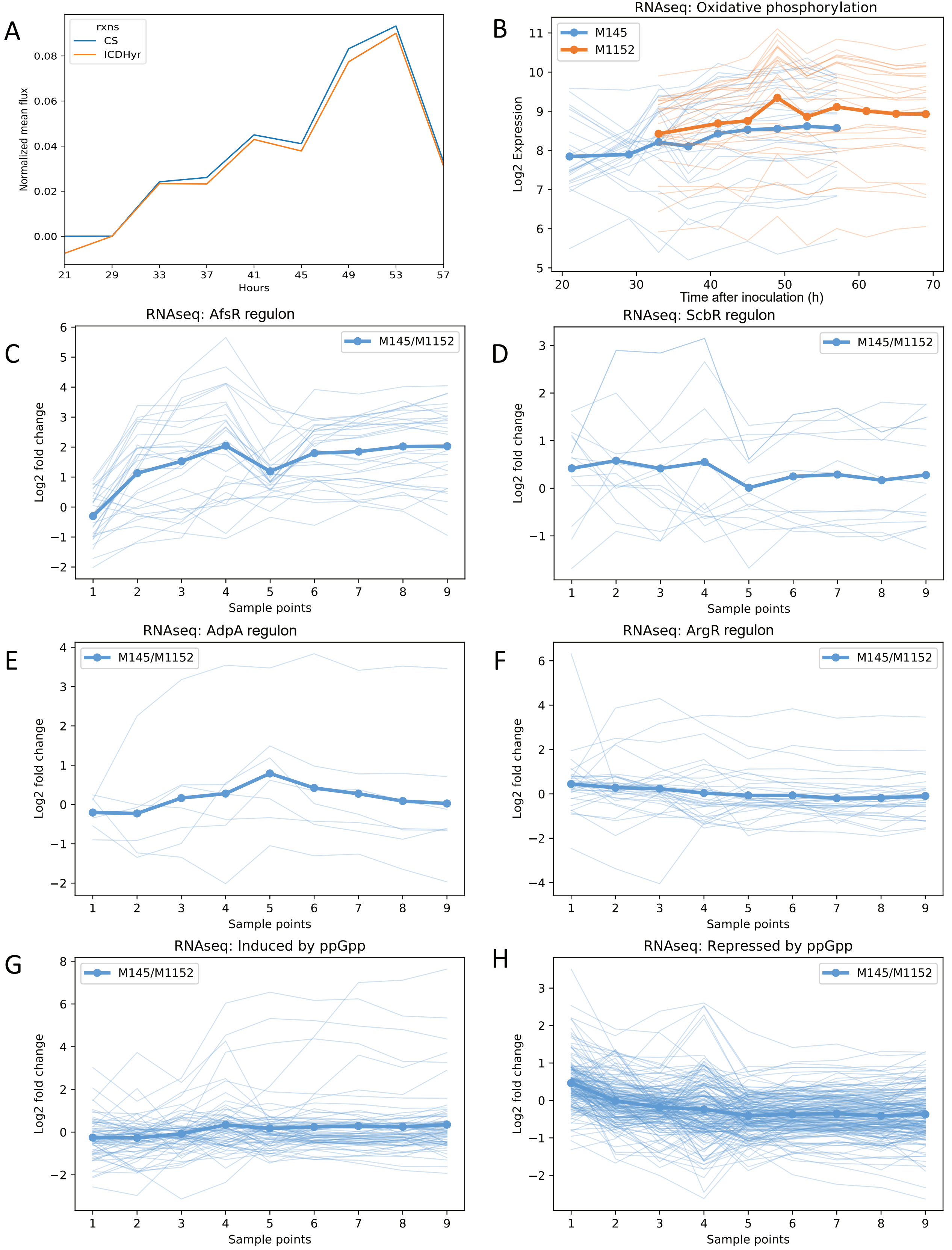

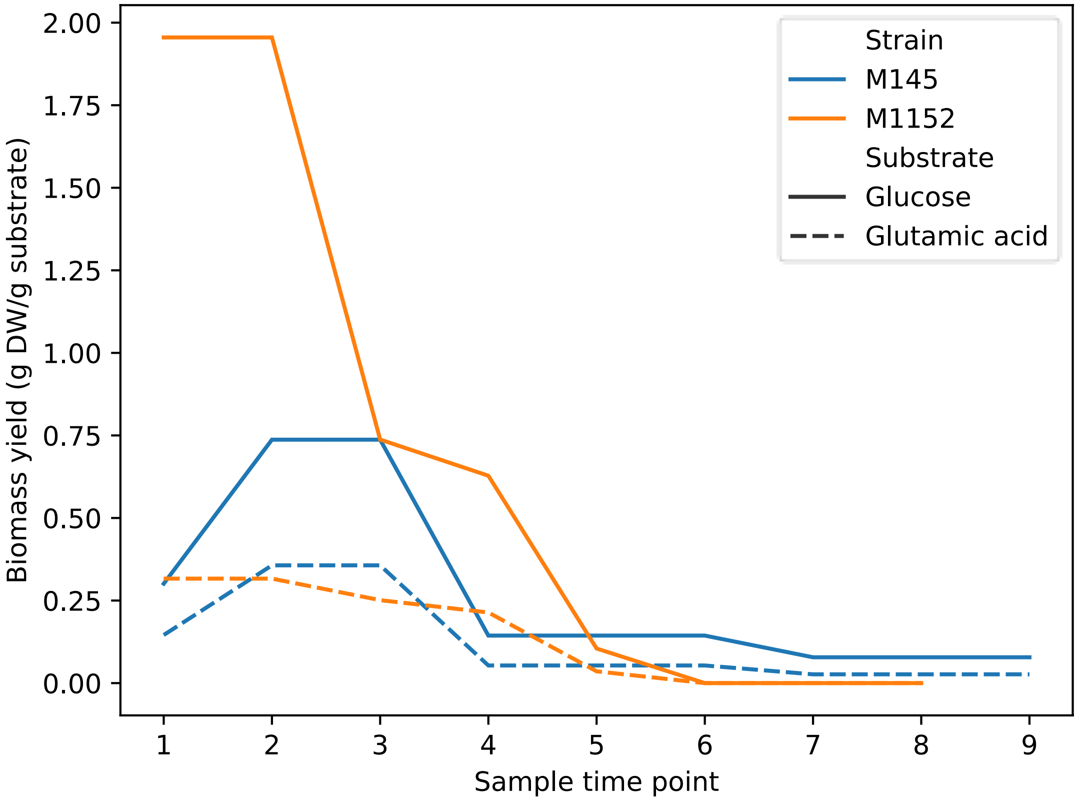

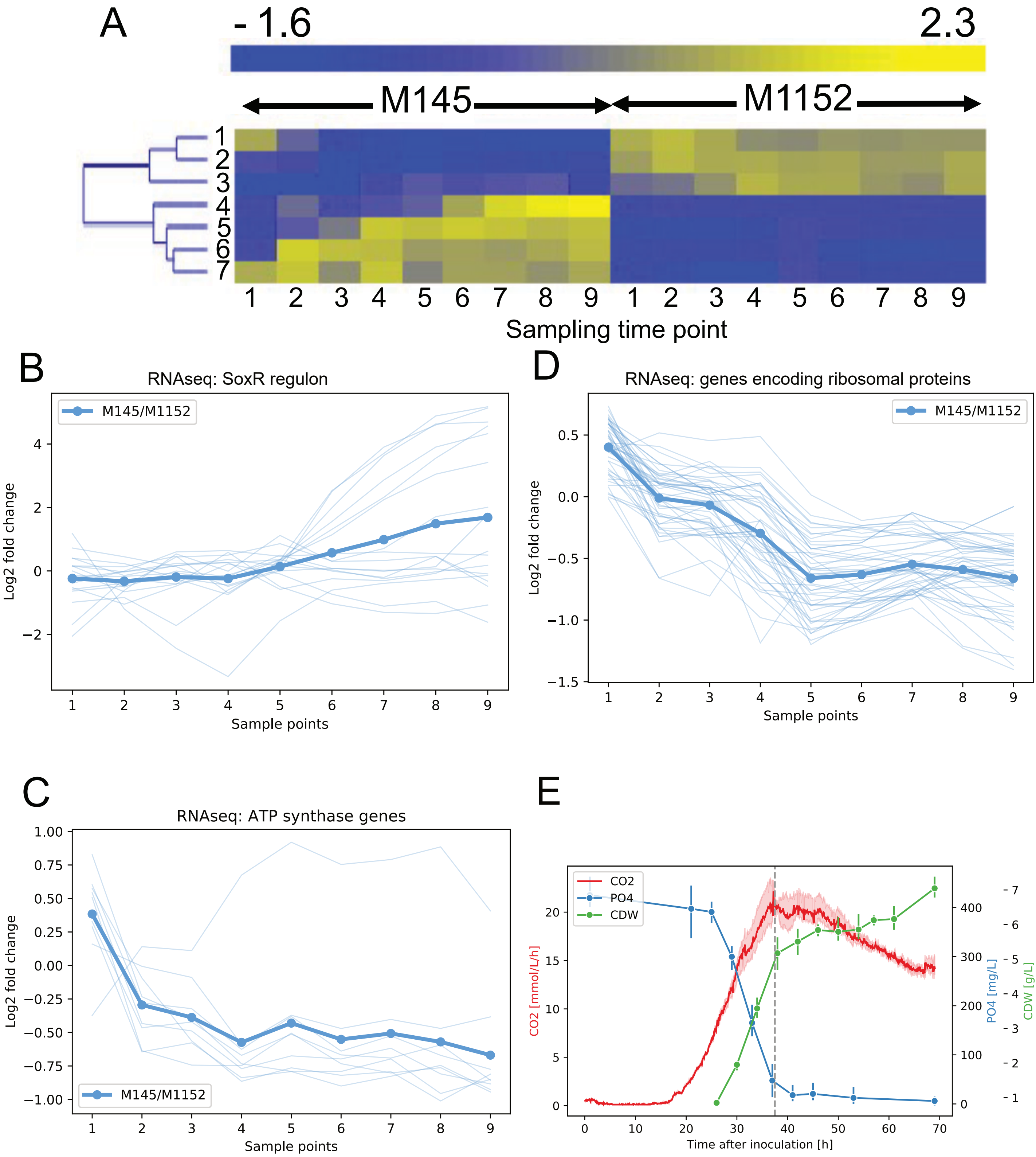

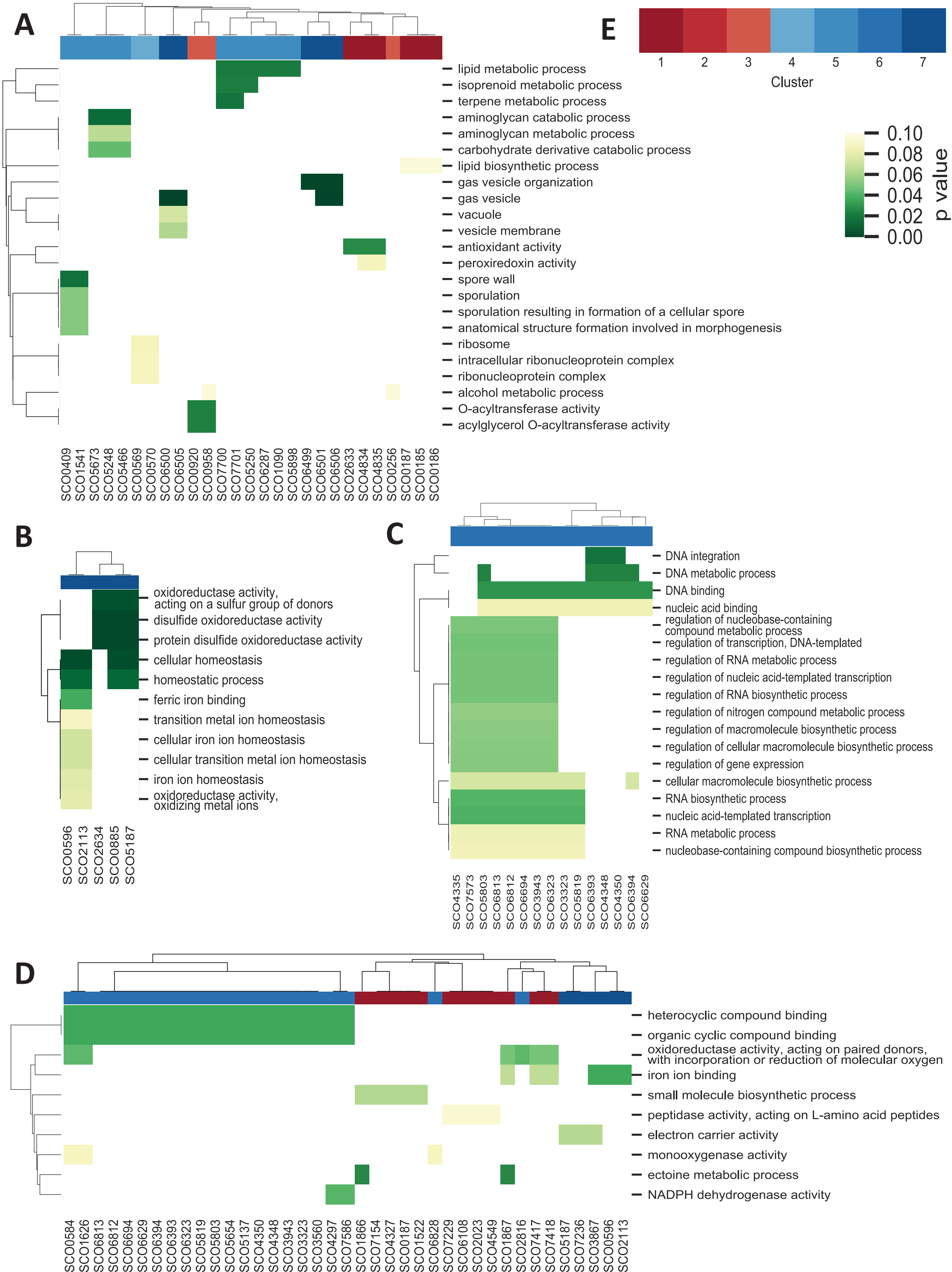

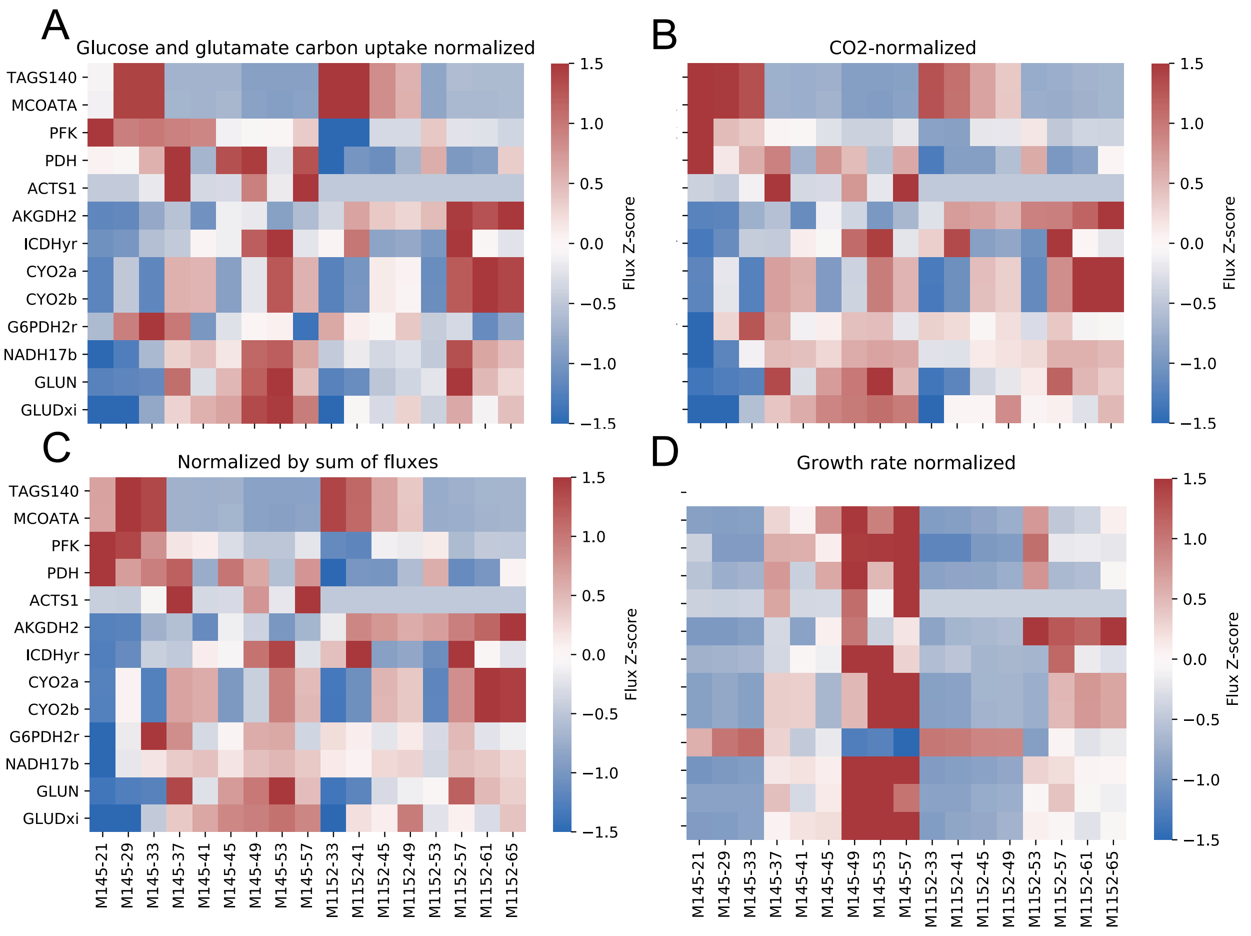
